# Modulation of gp130 Cytokine Receptor Family by Herpes Simplex Virus 1 as a Strategy for Immune Evasion

**DOI:** 10.64898/2026.10.06.757054

**Authors:** Kimberly Foster-Lemieur, Sreenath Muraleedharan Suma, Meenu Ajith, Maria Kalamvoki

## Abstract

HSV-1 is a neurotropic human pathogen that establishes lifelong latency in sensory neurons. Periodic reactivation causes mucocutaneous lesions and, in some cases, severe corneal and neurological disease. While HSV-1 utilizes numerous viral proteins to extensively reprogram host signaling pathways and evade antiviral responses, how it alters host cytokine receptor networks remains poorly understood. Here, we demonstrate that HSV-1 infection selectively downmodulates multiple members of the gp130 (glycoprotein 130) cytokine receptor family, including gp130, Oncostatin M receptor β (OSMR-β), and leukemia inhibitory factor receptor (LIFR), while sparing IL-6Ra and CNTFRa (Ciliary Neurotrophic Factor Receptor). Because gp130 is the central signaling component of the IL-6 cytokine family and mediates JAK/STAT3 activation, its elimination likely represents a viral strategy to suppress inflammatory and antiviral signaling. Consistent with this, HSV-1 does not induce IL-6 mRNA expression in productively infected cells, suggesting coordinated inhibition of both cytokine production and receptor availability. Mechanistically, we found that the viral RNase VHS is at least partially responsible for gp130 family receptor downmodulation, likely in combination with virus-induced endocytosis/degradation pathways. Functionally, activation of gp130 signaling significantly impairs HSV-1 infection, demonstrating that suppression of antiviral cytokine responses is critical for virus replication. Finally, we determined that CNTF causes reactivation of latent HSV-1 using the LUHMES *in vitro* latency model, illustrating CNTFRa signaling contribution to latency and reactivation. A similar effect has been reported for IL-6. Together, these findings reveal a previously unrecognized immune evasion mechanism by which HSV-1 remodels gp130 cytokine signaling to promote infection.

**Importance:** Successful infection requires HSV-1 to overcome innate immunity that would otherwise restrict viral replication and spread. However, how the virus alters cytokine receptor signaling at the cell surface remains poorly understood. The gp130 family of cytokine receptors represents an important signaling network that governs immunity. These receptors mediate responses to IL-6 family cytokines and regulate antiviral defense, inflammation, cell survival, and neuronal function. Because individual gp130 family receptors can initiate distinct responses, understanding how HSV-1 alters these receptors is important for defining how the virus reshapes host cytokine networks. Here, we report that HSV-1 downmodulates specific gp130 family members to evade host antiviral and inflammatory signaling. However, CNTFRa, a receptor vital to neuronal function is preserved. Characterizing this distinct layer of HSV-1 immune evasion broadens our understanding of virus–host interactions and highlights novel signaling pathways that can be targeted to maximize host antiviral defenses.

## Introduction

Herpes Simplex Virus 1 (HSV-1) is a highly prevalent human pathogen, with more than 70% of the global population harboring latent infections (1, 2). Primary infection occurs in mucosal epithelial cells, after which the virus establishes lifelong latency in sensory neurons (1). Reactivation most commonly causes orofacial lesions, but in immunocompromised individuals, recurrent infection can lead to severe disease, including encephalitis and keratitis (1, 3). The ability of HSV-1 to transition between lytic infection and latency is fundamental to lifelong persistence and depends on sophisticated mechanisms that evade host immune surveillance (1, 4). Accordingly, HSV-1 encodes several immune-modulatory proteins, including infected cell protein 0 (ICP0) and virion host shutoff (VHS), which suppress antiviral defenses and reshape host signaling pathways (1, 5). However, despite the central role of cytokine signaling in antiviral immunity, how HSV-1 manipulates host cytokine receptor networks remains poorly understood.

One of the earliest host responses to HSV-1 infection is activation of inflammatory cytokine signaling pathways. Infection of mucosal epithelial cells induces interleukin-6 (IL-6) production, triggering Janus kinase (JAK) activation and downstream phosphorylation of signal transducer and activator of transcription 3 (STAT3), a master regulator of acute inflammatory and antiviral transcriptional programs (6, 7). IL-6 signaling coordinates expression of interferon-stimulated genes (ISGs), inflammatory chemokines, and survival pathways that collectively restrict viral spread and recruit immune effector cells (8). Consequently, successful viral infection requires mechanisms to disrupt or redirect this signaling axis.

Central to this pathway is glycoprotein 130 (gp130), the shared signal-transducing receptor subunit for the IL-6 family of cytokine. gp130 integrates signaling from IL-6, IL-11, IL-31, leukemia inhibitory factor (LIF), ciliary neurotrophic factor (CNTF), oncostatin M (OSM), cardiotrophin-1 (CT-1), and related cytokines (9). Following ligand binding to the cognate α-receptor chain, gp130 undergoes homo- or heterodimerization, initiating JAK/STAT3, PI3K/AKT, and MAPK signaling cascades (9, 10). Because gp130 signaling simultaneously regulates inflammation, antiviral defense, cellular stress responses, proliferation, and survival, it represents a critical immunological control node during infection.

Importantly, gp130 signaling is uniquely positioned to amplify immune responses across tissues. Unlike many cytokine receptors, gp130 is ubiquitously expressed, enabling nearly all cell types to respond to IL-6 family cytokines. In addition to classical membrane receptor signaling, IL-6 can signal through soluble IL-6 receptor α (sIL-6Rα) generated by proteolytic shedding during inflammation, particularly by infiltrating neutrophils (11, 12). This “trans-signaling” mechanism allows IL-6/sIL-6Rα complexes to directly activate gp130 on neighboring cells, independent of surface IL-6Rα expression (11). As a result, gp130 functions as a broad inflammatory amplifier capable of rapidly propagating antiviral signaling throughout infected tissues (13, 14). Activation of this pathway induces STAT3- dependent transcription of antiviral and inflammatory genes, which are essential for limiting infection and coordinating immune responses (15, 16).

Given its central role in antiviral immunity, gp130 signaling has emerged as a frequent target of viral immune evasion. Multiple herpesviruses have evolved mechanisms to manipulate this pathway, underscoring its importance for viral persistence. Kaposi’s sarcoma-associated herpesvirus (KSHV) encodes viral IL-6 (vIL-6), which directly engages gp130 to promote immune evasion, cell survival, and viral pathogenesis (17, 18). Epstein–Barr virus (EBV) encodes a viral IL-10 homolog that modulates STAT3 activation, driving nasopharyngeal carcinoma (NPC) by regulating the JAK/STAT pathway to enhance tumor cell proliferation, migration, invasion, and survival (19, 20). Human cytomegalovirus (HCMV) similarly exploits IL-6/STAT3 signaling through the viral chemokine receptor US28 to induce IL-6 secretion and STAT3 activation, thereby promoting proliferation and persistence (21). Beyond upstream cytokines, many viruses directly target STAT3 itself, altering its activation or localization to suppress antiviral transcriptional programs (7). Collectively, these observations identify gp130/STAT3 signaling as a conserved and highly vulnerable hub exploited during herpesvirus infection.

Despite this, whether HSV-1 directly remodels gp130 receptor networks has not been investigated. Here, we show that HSV-1 selectively down-modulates multiple gp130 family receptor members, including gp130, OSMRβ, LIFR, and IL-31R, at both the mRNA and protein levels across immortalized and primary cell types relevant to natural infection. Deletion of VHS partially rescued receptor expression, implicating VHS in receptor down-modulation. Strikingly, this suppression was selective rather than global, as CNTFRα and IL-6Rα were preserved during infection.

The selective preservation of CNTFRα may be particularly significant in the context of viral latency. CNTF signaling through the CNTFRα–LIFR–gp130 complex is strongly neurotrophic and promotes neuronal survival (22), a process essential for maintenance of latent infection within sensory neurons. These observations raise the possibility that HSV-1 strategically rewires gp130 signaling to suppress inflammatory and antiviral STAT3 responses while preserving neuroprotective signaling pathways that support neuronal survival and long-term persistence. Mechanistically, we demonstrate that activation of the gp130 pathway inhibits virus infection. Stimulation with exogenous recombinant cytokine ligands significantly impaired viral growth, indicating that intact gp130 signaling restricts productive infection. Conversely, CNTF reactivated HSV-1 from latently infected cultures. Together, our findings support a model in which HSV-1 selectively remodels gp130 cytokine receptor networks to blunt antiviral immunity while preserving signaling pathways favorable for the latent stage of the virus. This work identifies gp130 receptor remodeling as a previously unrecognized immune evasion strategy linking cytokine signaling, neuronal biology, and herpesvirus persistence.

## Results

### HSV-1 selectively downregulates members of the gp130 receptor family

gp130 is a ubiquitously expressed, shared signal-transducing subunit that forms signaling complexes with multiple cytokine receptors and plays a key role in mediating inflammatory responses (9, 11). Formation of oligomeric signaling complexes with gp130 is the basis of classic signaling motifs within this receptor system; however, signaling within this family is diverse and widespread, reflecting the multitude of receptor combinations that can induce distinct downstream signaling cascades (23, 24). While HSV-1 has evolved sophisticated strategies to reprogram host signaling networks to evade antiviral immunity and sustain replication and persistence, the extent to which HSV-1 reshapes host cytokine receptor landscapes remains poorly defined (25-27).

Previous studies have implicated HSV-1 infection in altering IL-6 signaling (28). Here, we sought to determine how HSV-1 affects individual members of the gp130 receptor family. To characterize this effect, we performed a kinetic analysis in hTERT-HEL infected with HSV-1(F). Cells were infected at a high virus dose (10 PFU/cell) to ensure infection of all cells, and the expression of multiple gp130 receptor family members was examined over the course of infection. As shown in Figure 1A, gp130, IL-31R, OSMR-β, and LIFR were markedly downregulated. This effect was observed as soon as 5 h post-infection (hpi). Conversely, IL-6Ra and CNTFRa were not significantly affected (Figure 1A). TNFR1 served as a positive control because this cytokine receptor has previously been reported to be downregulated during HSV-1 infection (29). IL-31R appeared as two bands on the western blot, likely representing the oligomeric and monomeric forms (Figure 1A).

**Figure 1:**
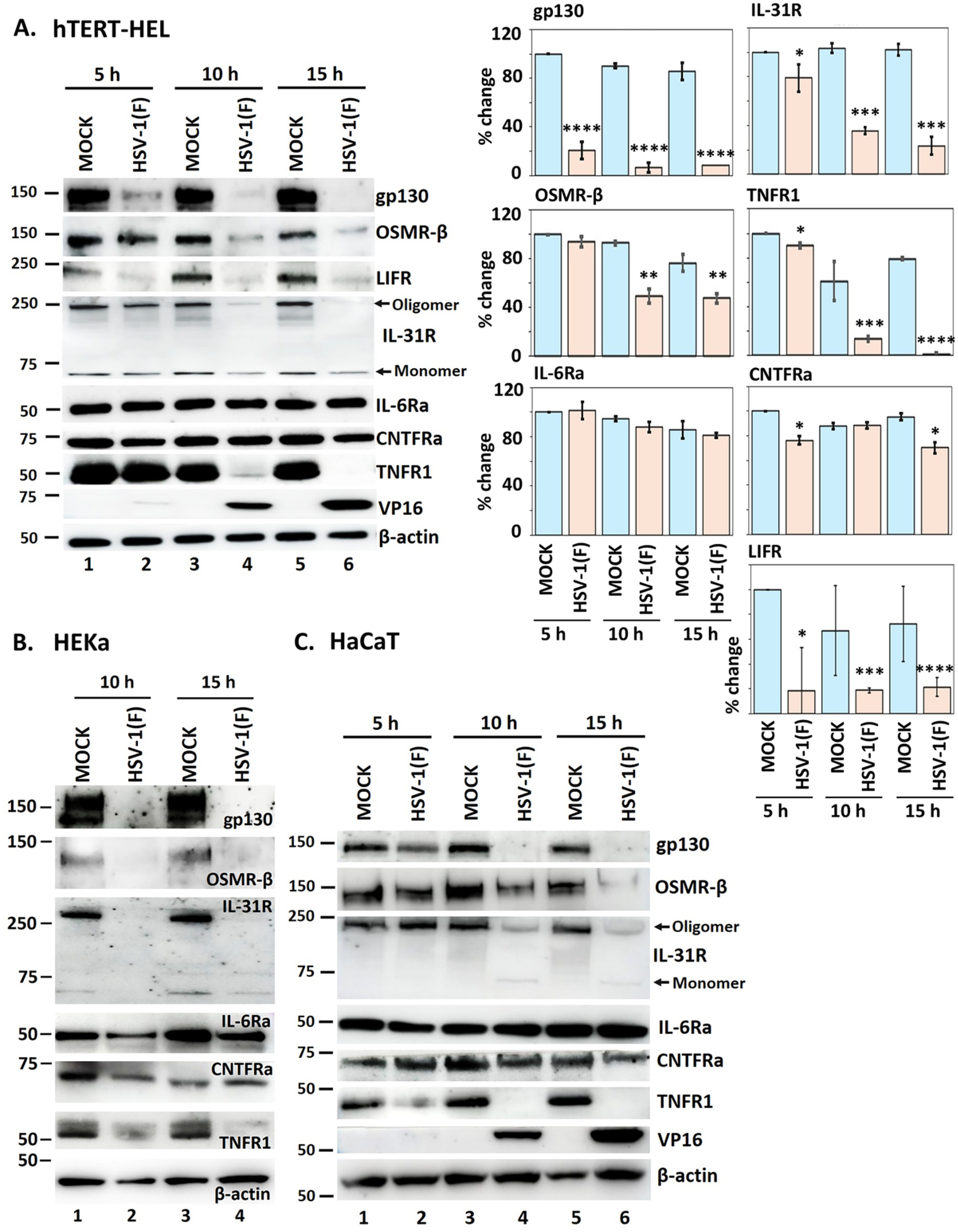
HSV-1 infection downmodulates selected members of the gp130 receptor family. **A.** hTERT-HEL cells were infected with HSV-1(F) (10 PFU/cell) or left uninfected. The cells were harvested at 5,10, and 15 hpi and equal amounts of proteins from total cell lysates were analyzed for gp130, OSMR-β, LIFR, IL-31R, IL-6Rα, CNTFRα, and TNFR1. VP16 served as a control for infection and β-actin as a loading control. Quantification of protein signal from three independent experiments was done using ImageJ. P<0.05*, P<0.01**, P<0.001***, P<0.0001**** with two-tailed Student’s *t*- test. **B.** Human primary epidermal keratinocytes (HEKa) were infected with HSV-1(F) (10 PFU/cell) or left uninfected. The cells were harvested at 10 and 15 hpi and equal amounts of proteins from total cell lysates were analyzed for gp130, OSMR-β, IL-31R, IL-6Rα, CNTFRa, and TNFR1. VP16 served as a control for infection and β-actin as a loading control. **C.** HaCaT cells (immortalized human keratinocytes) were infected with HSV-1(F) (10 PFU/cell) or left uninfected. The cells were harvested at 5, 10, and 15 hpi and equal amounts of proteins from total cell lysates were analyzed as before.

To determine whether this receptor remodeling was specific to the immortalized fibroblasts, we repeated this experiment in both immortalized (HaCaT) and primary human adult keratinocytes (HEKa). The same pattern of receptor downmodulation was observed in both cell types (Figure 1B-C), demonstrating that HSV-1-mediated downmodulation of gp130 family members is not restricted to a single cellular model. Protein downmodulation was quantified using ImageJ from three independent experiments (Figure 1A). Together, these data indicate that HSV-1 rapidly remodels the composition of gp130 family cytokine receptors during infection.

### The ICP0/CIN85 interaction is not required for gp130 receptor family internalization

Several viral proteins, including ICP0, modulate host defense mechanisms (1, 5, 30). We previously identified a viral strategy involving ICP0, which forms a complex with CIN85 (Cbl-interacting protein of 85 kDa), an endocytic adaptor protein, and accelerates removal of cell surface receptors. The ICP0– CIN85–Cbl complex promotes the internalization of EGFR and Nectin-1, the latter of which is a major HSV-1 entry receptor (31-34). This receptor internalization enhances viral spread by preventing virus entry to uninfected cells (32). However, whether this same mechanism contributes to the downmodulation of gp130 family receptors has not been investigated.

To determine whether the ICP0–CIN85 interaction contributes to gp130 family receptor downmodulation, we performed a kinetic analysis using HSV-1 (F) and the ICP0 Δ244–277 mutant virus (31). ICP0 Δ244–277 lacks two motifs required for CIN85 binding, thereby reducing the ICP0– CIN85 interaction and preventing ICP0 localization to CIN85 endosomes (31). We infected hTERT-HEL cells with the two viruses at 5 PFU/cell, and gp130 family receptor levels were examined at 5, 10, and 15 hpi. The gp130, IL-31R, and TNFR1 receptors were downregulated in cells infected with the ICP0 mutant, comparable to WT virus infections, whereas OSMR-β showed a small delay (Figure 2A). Together, these data indicate that gp130 family receptor downmodulation during HSV-1 infection is largely independent of the previously characterized ICP0–CIN85 endocytosis pathway.

**Figure 2:**
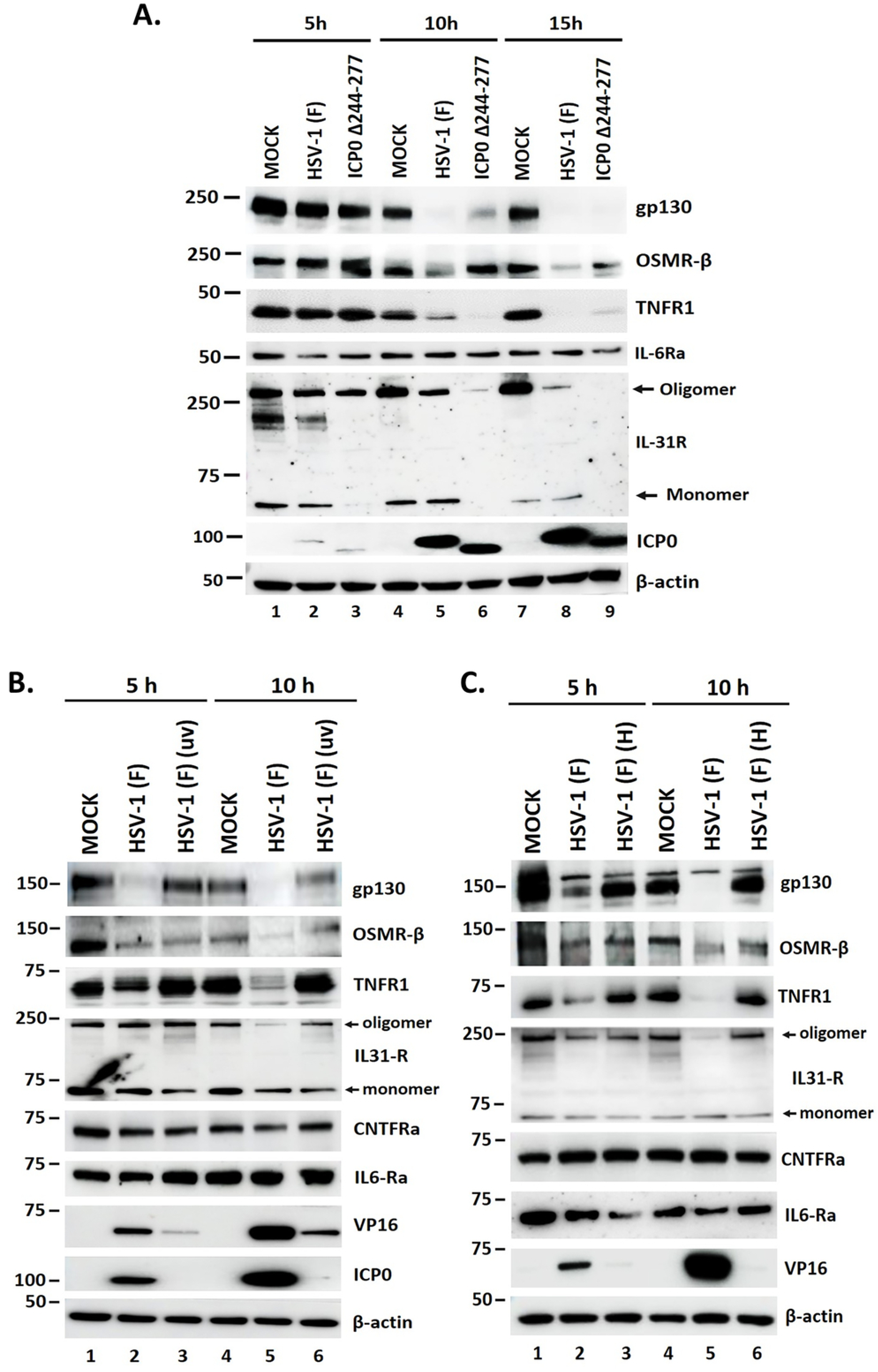
The ICP0/CIN85 interaction is dispensable for gp130 receptor family internalization, while UV-inactivated HSV-1 partially downmodulates certain family members. **A.** hTERT-HEL cells were infected with HSV-1(F) and ICP0 Δ244-277 virus (5 PFU/cell) or left uninfected. The cells were harvested at 5,10, and 15 hpi and equal amounts of proteins from total cell lysates were analyzed for gp130, OSMR-β, TNFR1, IL6-Rα, and IL-31R. ICP0 served as a control for infection and β-actin as a loading control. B. hTERT-HEL cells were infected with HSV-1(F) and uv-inactivated virus (20 PFU/cell) or left uninfected. The cells were harvested at 5 and 10 hpi and equal amounts of proteins from total cell lysates were analyzed for gp130, OSMR-β, TNFR1, IL6-Rα, and IL-31R. VP16 served as a control for infection and β-actin as a loading control. C. hTERT-HEL cells were infected with HSV-1(F) and heat-inactivated virus (20 PFU/cell) or left uninfected. The cells were harvested at 5 and 10 hpi and equal amounts of proteins from total cell lysates were analyzed as before.

To determine whether receptor downmodulation is triggered by viral entry or requires *de novo* viral gene expression, we next examined whether UV-inactivated HSV-1 could promote receptor loss. UV-inactivated virus was generated by exposing purified HSV-1 to 254-nm UV light, and the efficiency of virus inactivation was confirmed by plaque assays and testing for viral gene expression (Figure 2B). hTERT-HEL cells were infected with HSV-1(F) at 20 PFU/cell, exposed to an equivalent dose of UV-inactivated virus, or left uninfected. Cells were harvested at 5 and 10 hpi and analyzed for gp130 family receptors and viral gene expression. Partial downmodulation was observed for several receptors, including gp130, OSMR-β, and IL-31R, following exposure to UV-inactivated virus (Figure 2B). VP16 and ICP0 served as controls for UV inactivation. ICP0, an immediate-early protein that requires *de novo* expression, was not detected in cells exposed to UV-treated virus, whereas VP16, a tegument protein of the virus, was detected at low levels compared to *de novo* synthesis, which represent the amount present on the virion (Figure 2B). These findings indicate that receptor downmodulation occurs, at least in part, in the absence of *de novo* viral gene expression and therefore implicate virion-associated factors present at the time of entry. Additionally, we examined the impact of heat-inactivated virus, a treatment that disrupts virion-associated proteins and prevents productive infection. As expected, heat-inactivated virus did not induce receptor loss (Figure 2C).

Together, these data indicate that HSV-1 employs one or more virion-associated proteins to initiate gp130 family receptor downmodulation early during infection, while additional viral mechanisms may contribute at later stages.

### Reduction of gp130 receptor family transcripts occurs early during infection and involves the viral RNase, VHS

Our UV-inactivation experiments prompted us to investigate early events that may contribute to receptor loss, including changes at the transcript level. hTERT-HEL cells were infected with HSV-1(F) (10 PFU/cell), or left uninfected, and total RNA was extracted at 1, 3, and 10 hpi. Receptor transcript abundance was assessed by RT-qPCR. Reduced transcript levels were observed for gp130, OSMR-β, LIFR, and TNFR1, as well as for the cytokine IL-6. These reductions were apparent as early as 1 hpi, became more pronounced by 3 hpi, and by 10 hpi transcript levels were nearly undetectable (Figure 3A). Interestingly, CNTFRa exhibited the opposite pattern; its transcript levels increased significantly, with notable upregulation observed by 10 hpi (Figure 3A). This increase coincides with the sparing of CNTFRa observed at the protein level, suggesting that CNTFRa may contribute to HSV-1 infection.

**Figure 3:**
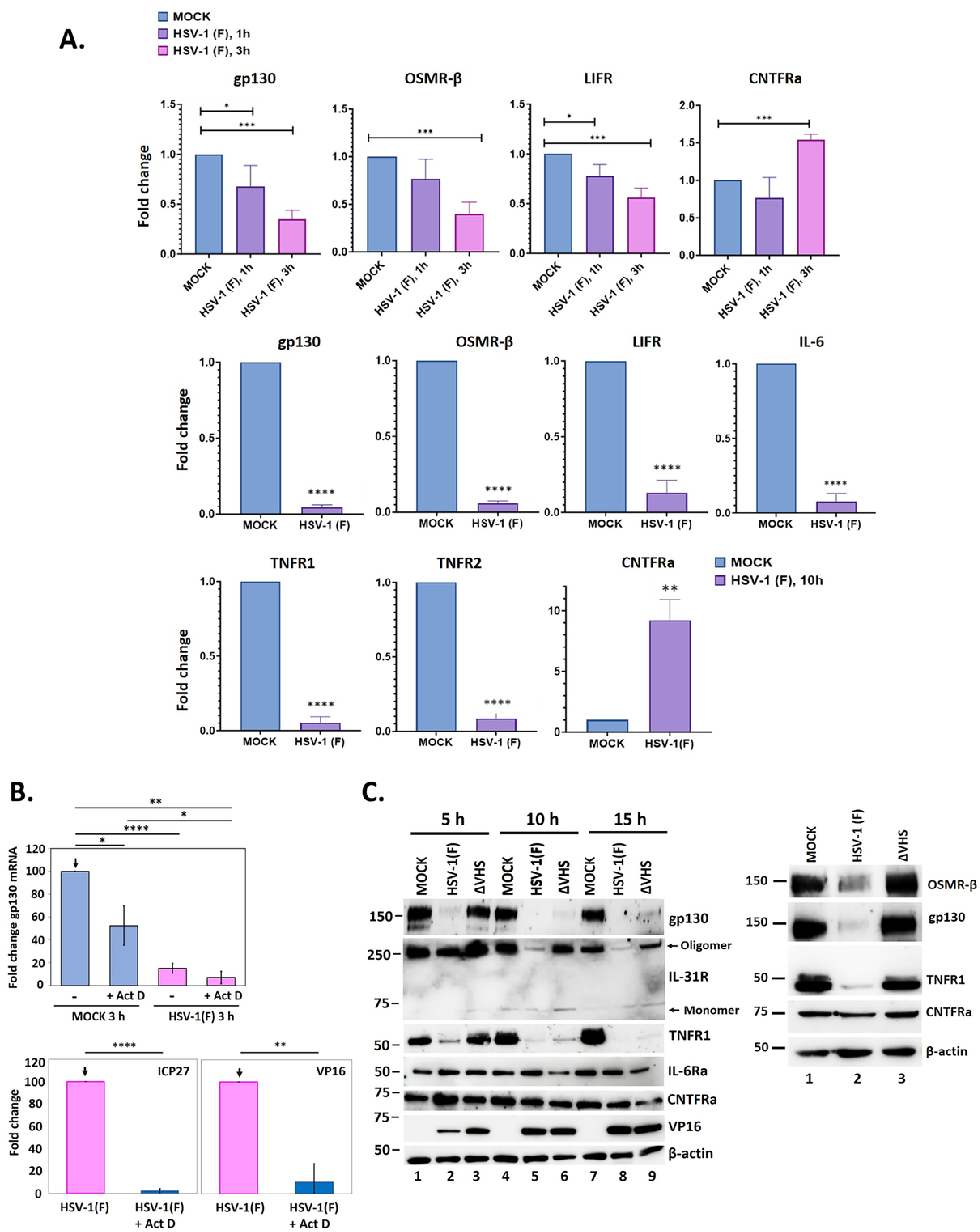
Rapid reduction in transcript levels of gp130 receptors, mediated, in part, by the HSV- 1 VHS. **A.** hTERT-HEL cells were infected with HSV-1(F) (10 PFU/cell) or remained uninfected. The cells were harvested at 1, 3, and 10 hpi, total RNA was extracted, and gp130 receptor and IL-6 cytokine gene expression was assessed by RT-qPCR. P < 0.05*, P < 0.01**, P < 0.001***, P < 0.0001****. **B.** hTERT-HEL cells were infected with HSV-1(F) (20 PFU/cell) or left uninfected. At 1 hpi, Act D (10 µM) was added to requisite samples, followed by collection in Trizol at 3 hpi. Total RNA was extracted, and gp130 expression was assessed by RT-qPCR. P < 0.05*, P < 0.01**, P < 0.001***, P < 0.0001****. **C.** hTERT-HEL cells were infected with HSV-1(F) and ΔVHS virus (10 PFU/cell), or left uninfected. The cells were harvested at 5, 10, and 15 hpi and equal amounts of proteins from total cell lysates were analyzed for gp130, OSMR-β, IL-31R, TNFR1, IL**-**6Rα, and CNTFRα. VP16 served as a control for infection and β-actin as a loading control. In addition, hTERT-HEL cells were also infected with HSV-1(F) or ΔVHS virus at low PFU (0.5 PFU/cell), and cell lysates were collected at 10 hpi and analyzed OSMR-β, gp130, TNFR1, and CNTFRa. β-actin served as a loading control.

To investigate the mechanism underlying this transcriptional reduction and whether it reflects accelerated mRNA decay, hTERT-HEL were either mock-infected or infected with HSV-1 (F) (20 PFU/cell). At 1 hpi, cells were treated with Act D (10 µM) or left untreated, and total RNA was extracted at 3 hpi. The transcript levels of gp130, ICP27, and VP16 were subsequently quantified by RT-qPCR. We observed that gp130 transcript levels significantly decreased within 3h in uninfected cells treated with Act D, reflecting the short transcript half-life (Figure 3B). In WT virus-infected cells, this decrease was greater than transcriptional inhibition alone, indicating active transcript decay (Figure 3B). While the addition of Act D did not change the impact of the virus on gp130 transcript levels, viral infection itself was significantly impaired, as indicated by reduced levels of viral transcripts (Figure 3B). The rapid loss of receptor transcripts, in the absence of viral gene expression, suggested the involvement of an early viral mechanism capable of targeting host mRNAs.

HSV-1 induces widespread degradation of host mRNAs through the activity of VHS, a viral RNase that promotes host transcript degradation and contributes to the suppression of antiviral responses (1, 35, 36). Because VHS is packaged within the virion tegument, it is delivered into the host cell upon viral entry and can promote host mRNA degradation during the first minutes of infection, prior to *de novo* viral gene expression (1, 32). VHS initiates cleavage of host mRNAs and cooperates with cellular mRNA decay machinery to promote their subsequent degradation (1). To test whether VHS has a role in gp130 downmodulation, we performed a kinetic experiment using a VHS-deficient virus (ΔVHS). Cells were infected with either WT or ΔVHS (10 PFU/cell) and equal amounts of proteins from total cell lysates were analyzed at 5, 10, and 15 hpi alongside uninfected controls. As expected, WT virus infection resulted in progressive loss of several gp130 family receptors. In contrast, infection with ΔVHS resulted in an early, but transient, rescue of expression for some of these receptors, including gp130, IL-31R. At later time-points, however, this rescue was diminished, with receptor levels trending toward those observed during WT virus infection. CNTFRa and IL-6Ra, which were resistant to downmodulation during WT infection, remained unaffected by the loss of VHS activity (Figure 3C). An additional analysis that was performed using lower virus dose yielded similar results (Figure 3C). Together, these data indicate that VHS contributes to gp130 family receptor downmodulation during the early stages of HSV-1 infection. However, the VHS-dependent phenotype was not sustained throughout the infection. Thus, while VHS likely contributes to early receptor modulation, these findings suggest that receptor downmodulation is not solely VHS-dependent and that additional viral and/or host mechanisms contribute as infection progresses.

### Downmodulation of gp130 family members occurs, at least in part, by lysosomal degradation

Here, we investigated the additional mechanisms driving the downregulation of this receptor family, as well as the intracellular fate of these receptors post-internalization. Typically, internalized receptors are degraded through the lysosomal pathway. Therefore, we analyzed a panel of inhibitors, including lysosomal and proteasomal inhibitors for their effect on gp130 family receptors. hTERT-HEL cells were infected with HSV-1 (F) (10 PFU/cell), followed by treatment at 1 hpi with either MG132 (15 µM), bafilomycin (BafA1) (15 µM), or CHX (100 µg/mL). Equal amounts of proteins from total cell lysates were then analyzed by western blot at 10 hpi (Figure 4A). We found that treatment with the proteasome inhibitor (MG132) caused a partial rescue of all receptors tested but also caused a severe delay in viral gene expression, which may account for the rescue. Whereas addition of the lysosomal/autophagy inhibitor (BafA1) significantly rescued all receptors and further increased the basal levels of gp130 and TNFR1 in uninfected cells, implying that these receptors have a natural turnover through the lysosomal pathway. BafA1, in contrast to MG132, caused only a small delay in viral gene expression. Finally, treatment with CHX resulted in a complete absence of gp130 and TNFR1 in both uninfected and infected conditions. The disappearance suggests that these receptors rely on continuous, *de novo* translation to maintain their basal levels, reflecting the short half-life of these proteins (Figure 4A). Regarding IL-31R, CHX treatment resulted in no change in uninfected conditions, but partially rescued receptor loss during infection, demonstrating that: 1) This receptor has a longer half-life than gp130 and TNFR1; and 2) because blocking translation resulted in observable rescue, this receptor is likely dependent on *de novo* viral protein synthesis for its clearance from the cell surface. These treatments left both IL6-Ra and CNTFRa relatively unaltered (Figure 4A). Together, these data indicate that lysosomal activity degrades some gp130 family members and TNFR1.

**Figure 4:**
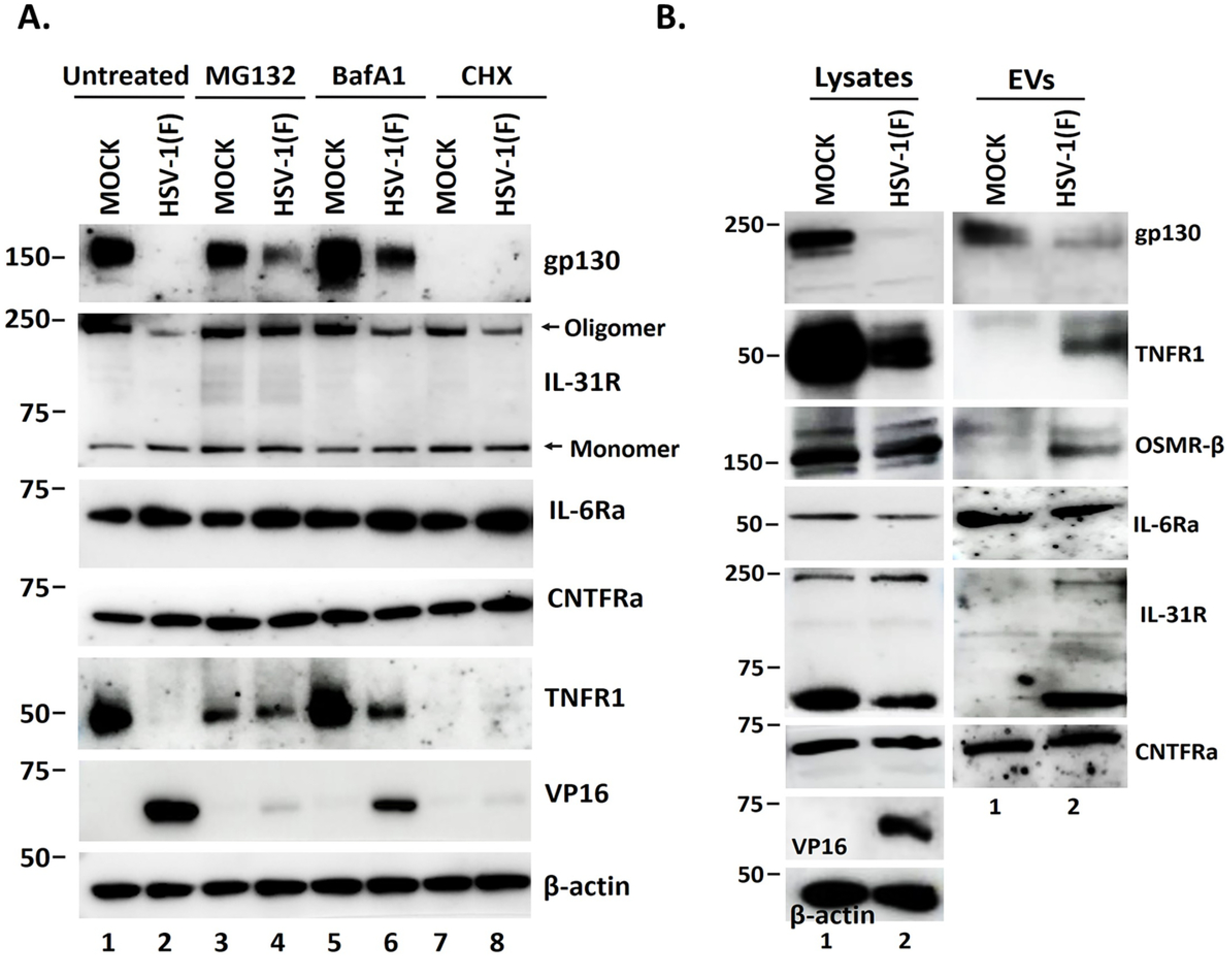
Mechanisms of gp130 receptor loss: role of proteasomes, lysosomes, and EVs. **A.** hTERT-HEL were uninfected or infected with HSV-1 (F) (10 PFU/cell). At 1 hpi, the cells were treated with MG132 (15 µM), bafilomycin (BafA1) (15 µM), and cycloheximide (CHX) (100 µg/ml). Cell lysates were collected at 10 hpi and analyzed via western blot for gp130, IL-31R, IL-6R, CNTFRa, and TNFR1. β-actin served as a loading control and VP16 as a control for infection. **B.** hTERT-HEL were infected with HSV-1 (F) (0.5 PFU/cell) or left uninfected. Culture supernatants were collected at 16 hpi. EVs were isolated from the supernatant after pelleting at 100,000 × *g* and analyzed by western blot for gp130, TNFR1, OSMRβ, IL6-Ra, IL-31R, and CNTFRa. β-actin served as a loading control and VP16 as a control for infection.

### gp130 family members enter exocytosis pathways following endocytosis

Typically, internalized receptors are directed to recycling and degradation pathways, or for exocytosis (34). In HSV-1-infected cells, several endosomal pathways divert cargo for exocytosis, as this represents a viral-immunoevasion strategy (31, 34, 37, 38). Here, we investigated whether gp130 family receptors are exocytosed through EVs. hTERT-HEL cells were infected with HSV-1 (F) (0.5 PFU/cell), or left uninfected, and EVs were collected from culture supernatants at 16 hpi, as we have previously described (39). Briefly, supernatants were filtered through 450 nm filters and concentrated using 100 kDa cutoff filter devices. EVs were pelleted by centrifugation at 100,000× *g,* and equal amounts of proteins were analyzed by western blot for gp130, TNFR-1, OSMR-β, IL6-Ra, IL-31R, and CNTFRa. For comparison, equal amounts of proteins from total cell lysates were also analyzed for these same receptors. β-actin was used as a loading control and VP16 as a control for infection. We observed that all tested receptors were at least partially present in EVs (Figure 4B).

Taken together, our data suggests that the selective clearance of gp130 family receptors involves early viral nuclease activity that clears the mRNA supply, alongside an endocytic pathway that transfers these receptors into lysosomes for degradation or for exocytosis through EVs.

### Pretreatment with sIL-6 restricts HSV-1 replication, but the virus efficiently counteracts this effect

STAT3 is a major signal transduction molecule downstream of gp130 family receptors involved in inflammatory responses (15). As such, efforts by pathogens to control this arm of immunity is widespread among herpesviruses and many other virus families, in both pro-viral and anti-viral contexts (40, 41). Therefore, we next sought to determine if HSV-1 infection functionally impairs gp130/JAK/STAT signaling, hypothesizing that activation of this pathway might inhibit HSV-1 replication.

To stimulate the gp130/JAK/STAT pathway, we used IL-6 bound to soluble IL-6Ra (rhIL-6/sIL-6Ra). This complex potently activates gp130 signaling independently of membrane-bound IL-6R expression on target cells (12). To assess pathway significance during infection, hTERT-HEL cells were pretreated with rhIL-6/sIL-6Ra for 16 h and then infected with HSV-1 (F) (10 PFU/cell) for 10 h. We observed STAT3 phosphorylation at tyrosine 705 after rhIL-6/sIL-6Ra treatment, indicative of pathway activation. Activation of the gp130/JAK/STAT pathway restricted infection, leading to significant delay in late viral gene expression (Figure 5A). These findings demonstrate that this pathway functions as a critical, early host immune response to control viral infection.

**Figure 5:**
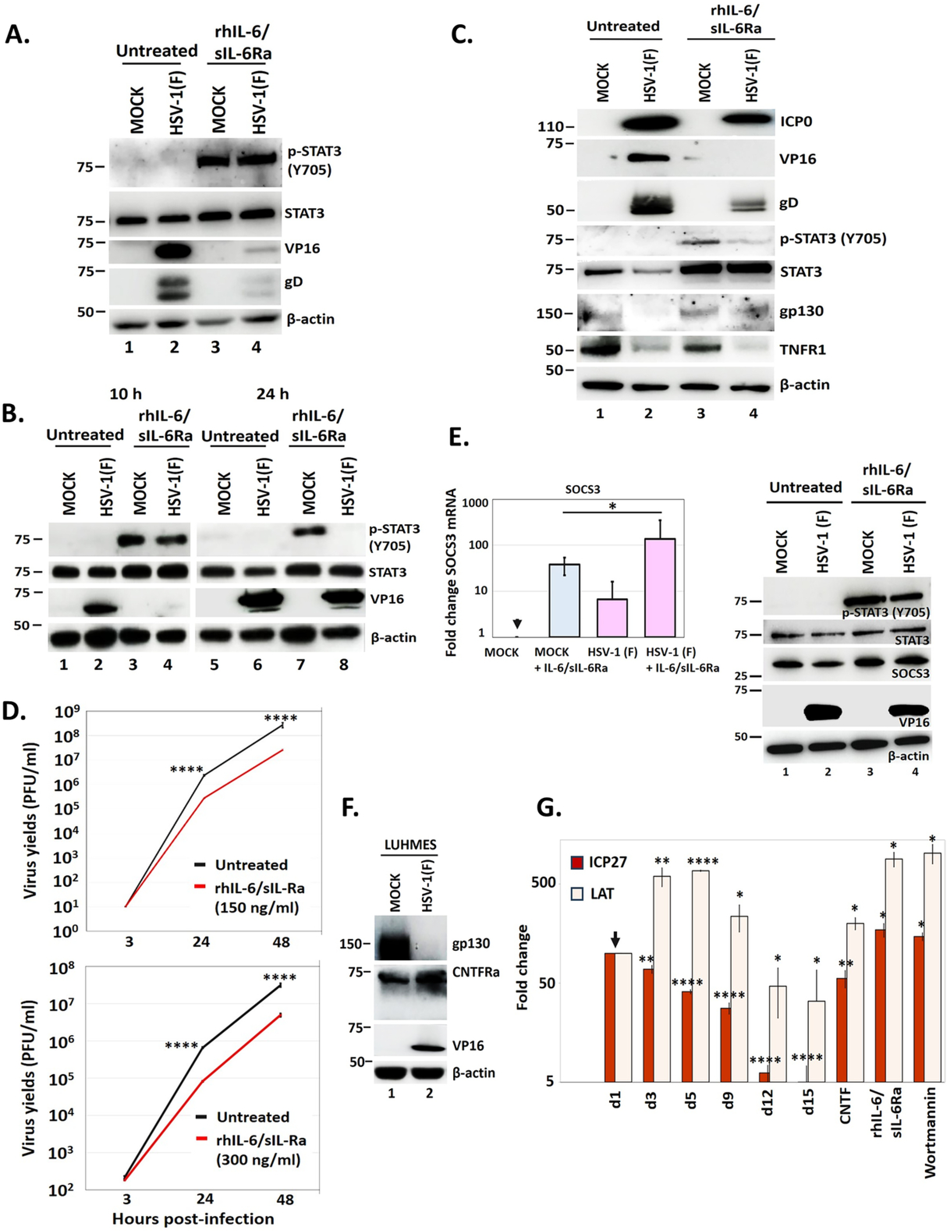
Growth defects of HSV-1 after treatment with rhIL-6/sIL6-Rα. **A.** hTERT-HEL cells were pretreated with rhIL-6/sIL-6Rα (300ng/ml) for 16 h or left untreated. Cells were then infected with HSV-1(F) (10 PFU/cell) or left uninfected. Cells were collected at 10 hpi and equal amounts of proteins from total cell lysates were assessed for p-STAT3 (Y705), total STAT3, viral proteins VP16 and gD, and for β-actin as a loading control. **B.** Same condition as in panel 5A, but cell lysates were collected at 10 and 24 hpi. **C.** Human primary epidermal keratinocytes (HEKa) were pretreated with rhIL-6/sIL -6Rα and infected with HSV-1(F), as in panels 5A-B. Cells were harvested at 10 hpi and equal amounts of proteins from total cell lysates were analyzed for the viral proteins ICP0, VP16, gD, and for p-STAT3 (Y705), total STAT3, gp130, TNFR1 with β-actin serving as a loading control. **D.** hTERT-HEL cells were pretreated with rhIL-6/sIL -6Rα (300 ng/ml) for 16 h or left untreated. Cells were then infected with HSV-1(F) (0.01 PFU/cell), harvested at 3, 24, and 48 hpi, and progeny virus production was quantified in Vero cells by plaque assays. **E.** hTERT-HEL cells were pretreated with rhIL-6/sIL -6Rα (300 ng/ml) for 16 h or left untreated. Cells were then infected with HSV-1(F) (0.1 PFU/cell) and harvested in Trizol at 6 hpi for transcript analysis. SOCS3 levels were assessed by RT-qPCR. P < 0.05*. In a parallel experiment, hTERT-HEL cells were pretreated with rhIL-6/sIL -6Rα (200 ng/ml) for 16 h or left untreated. These cells were then infected with HSV-1(F) (10 PFU/cell) and harvested at 10 hpi. Equal amounts of proteins from total cell lysates were analyzed for p-STAT3 (Y705), STAT3, SOCS3, viral protein VP16, and for β-actin as a loading control **F.** LUHMES cells were differentiated into mature dopamine-like neurons by exposure to tetracycline, GDNF, and dibutyryl-cAMP for 10 d. Cells were infected with HSV-1 (F) (10 PFU/cell), harvested 24 hpi, and equal amounts of proteins from total cell lysates were analyzed for gp130 and CNTFRα. VP16 served as a control for infection and β-actin as a loading control. **G.** LUHMES cells were differentiated into mature dopamine-like neurons by exposure to tetracycline, GDNF, and dibutyryl-cAMP for 10 d, using established protocol. The cultures were then infected with HSV-1 (F) (3 PFU/cell) in the presence of ACV. ACV was removed 48 h post-treatment, wherein an acute phase latency was established, characterized by high levels of latency associated transcript (LAT) and low levels of lytic gene expression such as ICP27 and VP16. At 19 dpi the cells were treated with CNTF (200 ng/ml), rhIL-6/sIL-6Rα (200 ng/ml), or wortmannin (PI3K kinase inhibitor, a positive control for reactivation, 2 μM). Samples were harvested on days 1, 3, 5, 9, 12, 15 and then 48 h after virus reactivation (day 21), and were analyzed for viral gene transcription by RT-qPCR. P < 0.05*, P < 0.01**, P < 0.001***, P < 0.0001****.

Viruses have evolved mechanisms to efficiently counteract host attempts to block viral replication and protein synthesis (3, 42). To this effect, we observed that HSV-1 efficiently inhibits IL-6/sIL-6Ra-activated STAT3 signaling. As the infection progressed, we found that HSV-1-infected cells exhibited reduced levels of p-STAT3 (Y705) compared with uninfected controls. Notably, phosphorylated STAT3 became undetectable by 24 hpi, despite total STAT3 expression remaining unchanged (Figure 5B). Viral gene expression was also enabled after the virus counteracted this antiviral signaling (Figure 5B). This phenotype was recapitulated in human primary keratinocytes, where HSV-1(F) similarly inhibited IL-6/sIL-Ra-induced p-STAT3 (Tyr702) (Figure 5C) to enable viral gene expression.

To explain why the virus counteracts the gp130 receptor signaling, we further assessed the impact on infection by measuring viral growth after pathway activation. hTERT-HEL cells were pretreated with rhIL-6/sIL-6Ra (150 ng/ml or 300 ng/ml) for 16 h or left untreated, then were infected with HSV-1(F) (0.01 PFU/cell) for 3, 24, or 48 h and progeny virus production was quantified by plaque assays. We observed a substantial and statistically significant growth defect by 24 hpi, continuing through the 48 hpi, indicating that IL6-induced stimulation of this pathway delays HSV-1 infection (Figure 5D).

SOCS3 is a negative regulator of the gp130/JAK/STAT pathway, acting as a regulatory brake to finetune the inflammatory signaling (43-45). We hypothesized that SOCS3 may be activated under these conditions and therefore evaluated its expression at the transcript level. For this, hTERT-HEL cells were pretreated for 16 h with rh-IL-6/sIL-6Rα (200 ng/ml), then infected with HSV-1 (F) (0.1 PFU/cell), and cells were harvested at 6 hpi for SOCS3 transcript analysis. In uninfected cells, treatment with rh-IL-6/sIL-6Rα stimulated SOCS3 expression, mitigating inflammatory signaling (Figure 5E). HSV-1(F) infection similarly elevated SOCS3, likely to suppress weak inflammatory signaling; however in IL-6-treated cells, infection further stimulated SOCS3 expression to counteract the high inflammation (Figure 5E). We repeated the experiment at a higher virus dose to determine if the SOCS3 phenotype is sustained. hTERT-HEL cells were pretreated, as above, for 16 h with rh-IL-6/sIL-6Rα (200 ng/mL), infected with HSV-1 (F) (10 PFU/cell), and harvested at 10 hpi. Although the effect was modest, SOCS3 displayed a mild elevation in IL-6 treated samples during infection compared to untreated samples that remained relatively stable. Concurrently, p-STAT3 presence denoted pathway activation, which correlated with a decrease in VP16, consistent with prior data (Figure 5E). It is possible that SOCS3 upregulation is responsible for the subsequent disappearance of p-STAT3, observed by 24 hpi.

Overall, these findings demonstrate that host inflammatory responses driven through the IL6-gp130-JAK/STAT3 pathway actively blunt viral replication. In response, HSV-1 efficiently counters this defense mechanism in a dual manner: it suppresses cytokine-induced STAT3 phosphorylation and stimulates SOCS3 expression, likely acting in concert to evade inflammatory responses and maintain cellular persistence.

### CNTF and IL6 drive HSV-1 latently infected neuronal cells to reactivation

Latency is established in sensory neurons within the trigeminal ganglia, wherein the HSV-1 genome exists in episomal form, a stable circularized version that allows for lifelong persistence (1, 46, 47). In this state, most viral genes involved in the lytic cycle are silenced, and instead, the latency associated transcript, LAT, is transcribed as the major viral gene product (1). This non-coding RNA product acts to maintain the latent state and protect infected neurons from cell death (1, 48). In contrast, reactivation, triggered by multiple stressors (UV light, immune suppression, PI3K/AKT pathway disruption), causes the resumption of the lytic cycle and the production of new virions (1, 49-52). Interestingly, while most gp130 family members are down-modulated in HSV-1–infected cells, we found that CNTFRα was spared (and even upregulated at the transcript level), offering a unique avenue to study how this reshaping of the immune landscape may favor HSV-1 latency.

To test this, we used a pre-established *in-vitro* latency model based on LUHMES (Lund human mesencephalic) cells. This is a conditionally immortalized, human embryonic neural precursor cell line that can be differentiated into mature dopamine-like neurons using tetracycline, GDNF (glial cell line-derived neurotrophic factor), and dibutyryl-cAMP (53). We first confirmed that gp130 is downmodulated in infected LUHMES, whereas CNTFRa is preserved, like in the other cell types, as discussed before (Figure 4F). We then differentiated LUHMES for 10 days and infected with HSV-1 (F) at 3 PFU/cell, where the virus establishes latency in the presence of acyclovir (ACV, 100 µM), which was removed 48 hpi. Replicate cultures were harvested on different days to verify latency establishment. Latency was verified by decrease in transcripts found in the lytic cycle of the virus, such as *ICP27*, a transcript from an immediate-early gene, and increase in LAT. At 19 dpi, cells were treated with rhCNTF (200 ng/mL), rhIL-6/sIL-6Ra (200 ng/mL), or wortmannin (2µM) (53). Wortmannin served as a positive control to viral reactivation, while IL6 served as an additional control for ligand-induced reactivation, based on established literature (54, 55). Samples were harvested at 48 h after virus reactivation and analyzed for viral gene transcription by RT-qPCR. Treatment with rhCNTF triggered a significant increase in *ICP27* transcript. Similar reactivation profiles were observed with rhIl-6/sIL- 6R and wortmannin (positive controls), indicating that rhCNTF drives HSV-1 exit from latency (Figure 5G). These data suggest a mechanism whereby CNTF likely acts as an extracellular signal that promotes HSV-1 reactivation.

### Analysis of the surfaceoma of HSV-1 infected cells confirmed gp130 loss, and pointed to alterations in signaling, trafficking, and cytoskeleton, upon HSV-1 infection

To confirm the downmodulation of gp130 family receptors from the cell surface during HSV-1 infection and to further assess the mechanism of receptor loss, we performed surface protein biotinylation coupled by proteomics analysis. During biotinylation, cells are incubated with sulfo-NHS-biotin, a membrane impermeable form of biotin at 4 °C to prevent endocytosis. Biotin reacts with the primary amines on extracellular lysine residues, and biotinylated proteins are then captured with neutravidin beads (56). Triplicate hTERT-HEL samples were infected with HSV-1 (F) (10 PFU/cell) or left uninfected, followed by cell surface biotinylation at 12 hpi for 30 min at 4 °C. After samples were collected, both surface proteins and cell lysates were analyzed for gp130 receptors. As depicted in a representative replicate, there was loss of gp130, OSMR-β, CNTFRa, and TNFR1 in both cell surface and lysates (Figure 6A). After validating our findings of gp130 receptor loss through cell surface labeling, we performed liquid chromatography-tandem mass spectrometry (LC-MS/MS).

**Figure 6:**
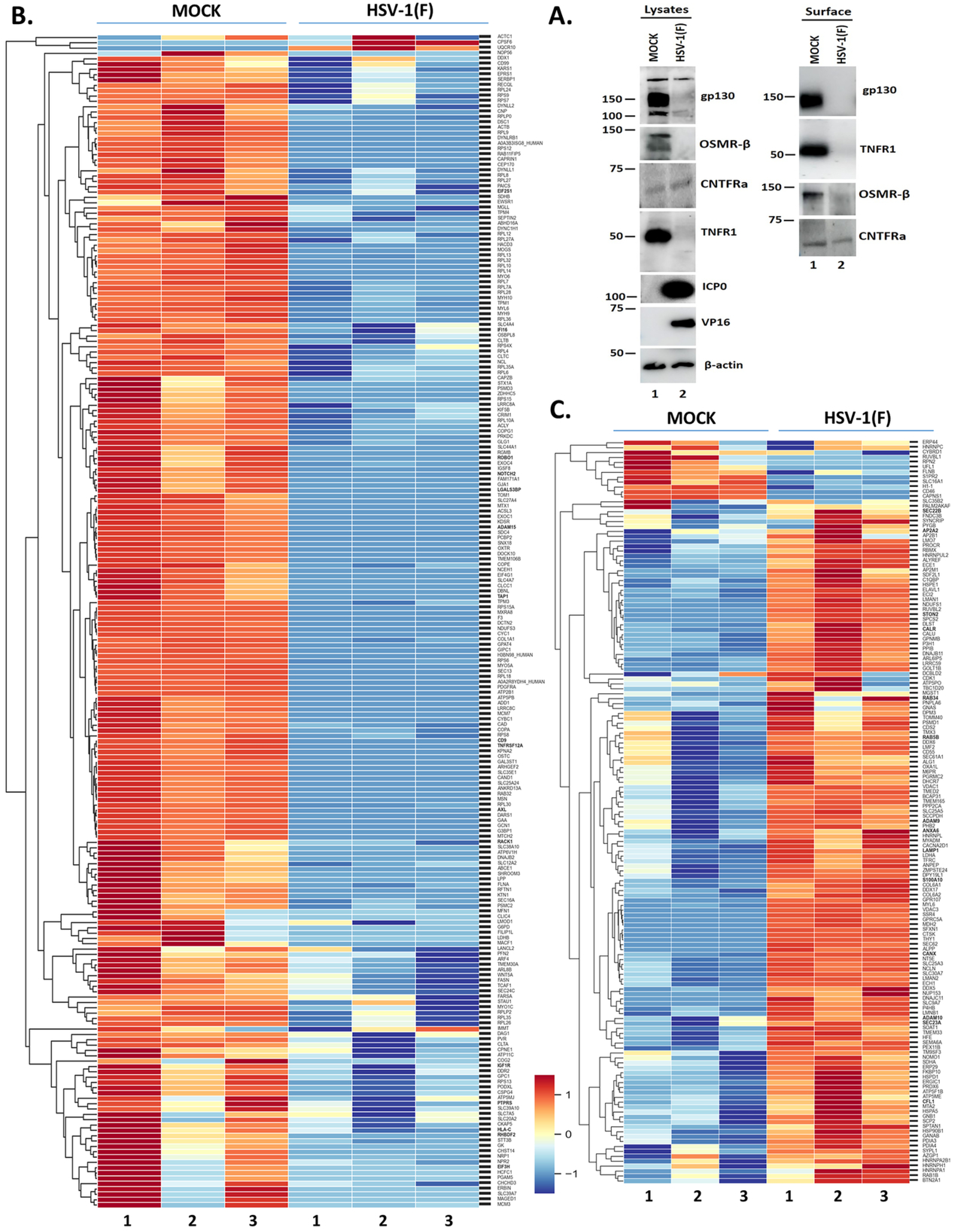
Analysis of surface proteins during HSV-1 infection. **A.** hTERT-HEL were infected with HSV-1 (F) (10/PFU/cell) or left uninfected. At 12 hpi, cells were biotinylated for 30 min. Labelled proteins and total cell lysates (equal amounts) were analyzed by western blot for gp130, OSMR-β, CNTFRα, and TNFR1. Viral proteins ICP0 and VP16 served as a control for infection and β-actin as a loading control. **B.** Heatmap depicting the most significantly downregulated proteins during HSV-1 infection (after surface biotinylation). Proteins related to gp130 signaling have been bolded. **C.** Heatmap depicting the most significantly upregulated proteins during HSV-1 infection (after surface biotinylation). Proteins potentially impacting gp130 signaling have been bolded.

Analysis for the most significantly upregulated and downregulated genes during infection revealed key targets within the gp130/JAK/STAT pathway. Infection was found to disrupt surface receptors, membrane organization, endocytosis, recycling, and trafficking. Key downmodulated targets included CD9, RHBDF2, and ADAM15. CD9 directly binds and stabilizes gp130 to promote downstream signaling, while RHBDF2 drives the surface maturation of ADAM17—the sheddase that cleaves membrane-bound IL-6Ra into trans-signaling-capable sIL-6Ra (Figure 6B) (57, 58). Furthermore, ADAM15 regulates membrane trafficking and fusion (59). By targeting this axis, the virus could destabilize gp130 and halt IL-6Ra proteolytic cleavage, effectively blunting pleiotropic signaling. Whereas, altered expression in CLTC, CLTA, FLNA, MSN, TOM1, ARF4, RAB11FIP5, RAB8A, and SEC24C indicates compromised receptor internalization, endosomal sorting, recycling, and plasma membrane delivery (Figure 6B) (60-64). Together, these changes could drive the loss of gp130-family receptors from the cell surface and reduce their replenishment, ultimately contributing to impaired JAK/STAT signaling.

In addition, HSV-1 elevates proteins associated with membrane proteolysis and vesicular/endolysosomal remodeling. ADAM10/ADAM9 upregulation suggests increased surface protease capacity, while AP2A1/AP2A2 and STON2 are known to support altered gp130 receptor internalization (Figures 6C) (65, 66). Increased levels of RAB5B/RAB34 and LAMP1 are consistent with enhanced endosomal/lysosomal trafficking, with ANXA6/S100A10 supporting membrane remodeling (Figures 6C) (66). These changes suggest mechanisms by which surface receptors could be internalized, redistributed, shed, or routed away from the plasma membrane during infection.

Overall, these data suggest an HSV-1-induced remodeling of the cell-surface–associated proteome, with clusters of proteins undergoing both up- and down-modulation in HSV-1(F)-infected cells relative to uninfected controls. This remodeling affects gp130 stability, cytoskeletal/membrane organization, receptor dynamics, and sorting/recycling pathways. Ultimately, this indicates a broad rewiring of structural and signaling components at the cell surface during infection, which alters surface dynamics to favor the infectious microenvironment and aid in viral persistence.

## Discussion

The IL-6 family of cytokines and the gp130 receptor system represent a critical innate immune defense mechanism through which the host counteracts viral pathogenesis. The pleiotropic nature of this cytokine family makes it an especially important target during viral dissemination within the host (67). Central to this signaling network is the ubiquitously expressed gp130 receptor, a shared signal-transducing subunit that, upon association with the appropriate α-receptor complex, activates the JAK/STAT pathway and drives the expression of inflammatory genes that coordinate innate immune responses (12, 68).

The pleiotropy of the IL-6 family is further reflected in the multiple mechanisms by which signaling can occur. In classic signaling, a cytokine binds its cognate α-receptor, followed by recruitment and dimerization of gp130, resulting in downstream signal transduction. In contrast, trans-signaling involves proteolytic cleavage of the membrane-bound α-receptor—most notably by ADAM17— followed by release of the soluble receptor (sIL-6Rα), which retains the ability to bind its ligand (69-71). The resulting cytokine–receptor complex can then engage gp130 directly on cells that do not express the membrane-bound α-receptor, thereby expanding the range of responsive cell types (71-74). Functionally, this mechanism broadens immune communication and can enhance host defense when a more robust inflammatory response is required. Conversely, gp130-mediated signaling can be negatively regulated by soluble gp130 (sgp130), which sequesters the cytokine–sIL-6R complex and prevents its interaction with membrane-bound gp130 (75, 76). In this capacity, sgp130 acts as a homeostatic regulator of IL-6 family signaling, helping to balance inflammatory responses and prevent excessive immune activation.

Multiple viruses exploit this pathway during infection, targeting components ranging from cell-surface receptors to downstream JAK/STAT signaling (7, 15). This suggests the existence of a conserved host defense mechanism that pathogens must overcome early on during infection. Accordingly, many viruses suppress critical innate immune responses by downregulating immune receptors at both the transcript and protein levels, as well as by inhibiting cytokine production (77, 78). These immune-evasion mechanisms have been observed across a broad spectrum of RNA and DNA viruses, yet their role in HSV-1 infection has remained largely unexplored (7, 79, 80).

Here, we demonstrate that similar principles apply to herpesviruses. Down-modulation of gp130 family receptors during HSV-1 infection likely represents an immunoevasion strategy. Manipulation of gp130 signaling appears to be a conserved strategy among herpesviruses. KSHV encodes vIL-6, which directly binds gp130 to promote immune evasion, cell survival, and viral pathogenesis (17, 18, 81-83). Although HSV-1 does not encode a cytokine homolog, our findings suggest that HSV-1 may achieve a functionally analogous outcome by selectively eliminating gp130 family receptors and thereby altering cytokine-mediated antiviral signaling.

To visually highlight these findings, we propose a model illustrating how HSV-1 modulates the gp130/JAK/STAT pathway to evade immune detection (Figure 7). Mechanistically, viral entry triggers a cascade of host-driven inflammatory and innate immune responses. Because this signaling pathway serves as a critical defense mechanism against incoming pathogens, the virus actively targets it to evade immune detection and maintain its lytic cycle. At the cell surface level, HSV-1 infection causes a selective downmodulation of key receptors, including the primary core signaling subunit, gp130.

**Figure 7:**
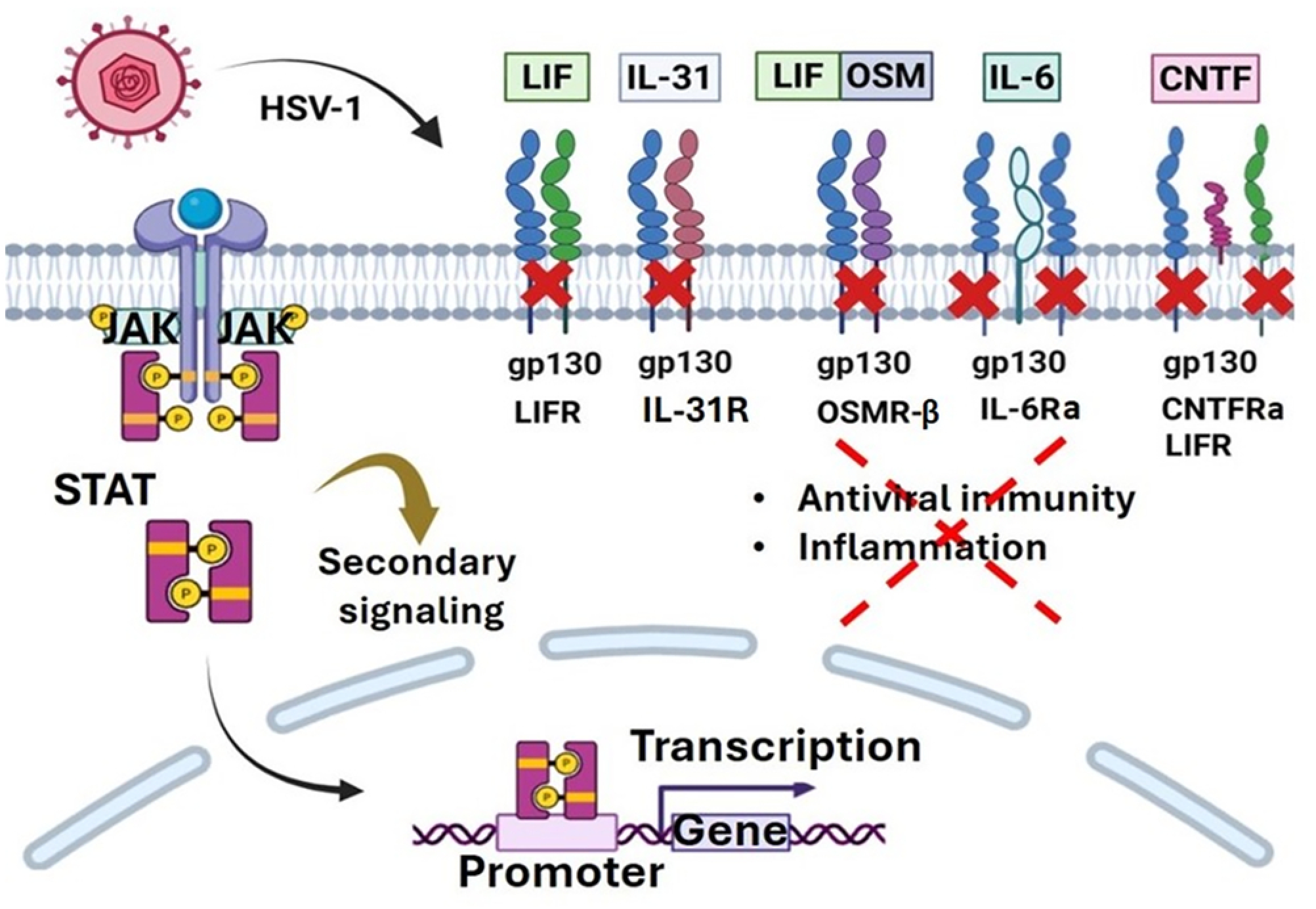
Model representing HSV-1 modulation of the gp130/JAK/STAT pathway. Viral entry triggers inflammatory responses including the production of IL6-family cytokines that signal through the gp130/JAK/STAT pathway. To counter host immune defenses, HSV-1 infection induces the selective downmodulation of gp130 family receptors, including the core signaling subunit, gp130. Other receptors removed from the surface include LIFR, OSMR-β, and IL-31R, whereas IL-6Ra and CNTFRa remain stable throughout infection. Downstream signaling, including STAT3 phosphorylation at Y705 is also abrogated.

Similarly, other gp130 family receptors are removed from the surface, whereas IL-6Ra and CNTFRa remain stable throughout the course of infection. The virus also affects downstream signaling, including STAT3 phosphorylation, thereby preventing inflammatory gene expression. Altogether, this mechanism represents a novel immune evasion strategy that has been underexplored yet exerts a major impact on the virus.

Our findings demonstrate that gp130, LIFR, OSMR-β, and IL-31R are significantly downregulated very early during infection, with decreases observed as soon as 3 hpi. This pattern of receptor downmodulation was consistent across multiple cell types supporting productive HSV-1 infection, indicating that HSV-1-induced downregulation of gp130 family members is not cell-type specific. We further analyzed transcript levels for the gp130 receptor family, which revealed rapid downmodulation of several receptors. Levels of gp130, OSMR-β, LIFR, TNFR1, and IL-6 transcripts decreased by 1 hpi, dropped further by 3 hpi, and became nearly undetectable by 10 hpi. Conversely, CNTFRa transcript levels increased, showing significant upregulation by 10 hpi. This increase matches the relative sparing of CNTFRa protein, suggesting a potential role for CNTFR in HSV-1 infection. These rapid declines suggested an early viral mechanism targets these host transcripts. When investigating the underlying viral mechanism, we identified overlapping strategies employed by the virus to suppress inflammatory responses through the gp130/JAK/STAT pathway. While this downmodulation operates largely independently of the previously characterized ICP0–CIN85 pathway, we discovered that the viral ribonuclease, VHS, plays a role early in infection. Cells infected with a VHS null virus exhibited a rescue of multiple receptors at early time points, consistent with the rapid nuclease activity characteristic of VHS. Also, UV-inactivated virus retained the ability to partially downmodulate these receptors indicating that this activity is associated with the virion. At later stages of infection, however, alternative mechanisms likely predominate as the ΔVHS phenotype does not persist.

We hypothesized that the pre-existing receptor pools might be targeted for degradation or exocytosis. Indeed, inhibiting lysosomal and autophagic pathways with BafA1 yielded a partial rescue. Also, a fraction of these receptors was detected in the extracellular space. Altogether, these data suggest that HSV-1 achieves selective clearance of gp130 family receptors through a coordinated, multi-pronged approach. Early VHS activity rapidly depletes the host mRNA pool, whereas a trafficking pathway shuttles these receptors into lysosomes for degradation and for exocytosis.

To validate the observed gp130 depletion from the cell-surface and expand our findings to other novel targets, we performed a complementary biotinylation capture of membrane proteins, followed by LC-MS/MS. This analysis revealed that multiple targets were significantly altered during infection, highlighting a global, virus-induced reorganization of the surface proteome. Specifically, the downregulation of genes involved in surface stability, proteolysis, endocytosis, sorting, and recycling points to additional mechanisms by which infection could remodel the gp130 receptor family, both by diminishing cell-surface presence and by disrupting the replenishment of receptors to the membrane. The reduction of RHBDF2 (iRhom2) and CD9 is particularly interesting; iRhom2 regulates ADAM17 trafficking, while CD9 can bind to and stabilize gp130 (57, 84, 85). Their diminished levels during infection suggest a multi-pronged viral strategy to target the gp130 receptor system.

Intriguingly, the upregulated genes also involve proteolysis, most notably, ADAM10. This membrane-bound metalloprotease is known to cleave the extracellular domain of membrane-bound gp130 to produce soluble sgp130, a negative regulator of gp130 signaling (86). Circulating sgp130 acts as an antagonist by binding to sIL-6R/IL-6 complexes, thereby blocking trans-signaling (87). Paradoxically, ADAM10 can also cleave IL-6Ra to produce sIL-6 and initiate the trans-signaling cascade; however, literature indicates that under acute phases of infection, increases in both IL-6 and sIL6-R may drive sgp130 production to dampen signaling (69, 86-89). Thus, an attractive hypothesis is that the infectious microenvironment drives the increase in cleavage through the gp130 arm, generating sgp130, either as a passive result of infection, or as a redundant mechanism to blunt the gp130/JAK/STAT pathway.

We also investigated the functional significance of the virus-induced downmodulation of selected gp130 family members and found it compromised viral gene expression. Upon ligand binding, gp130 family receptors activate kinases, including members of the Janus kinase family and phosphoinositide-3-kinases that convey signal to the nucleus, which result in inflammatory responses (11, 68). We discovered that pre-treatment with the recombinant human IL-6/IL-6Ra protein chimera activates these signaling pathways and this effect compromises viral gene expression. However, the virus counteracts this and eventually downmodulates the gp130 family receptors and reduces p-STAT3 (Y705). Additionally, we identified a secondary mechanism that suppresses this signaling axis via the upregulation of SOCS3. Upon evaluating transcript levels following pathway stimulation, we observed that viral infection induces SOCS3, a known negative regulator of the JAK/STAT3 pathway (90). This indicates that while the pathway itself is pleiotropically activated, the virus utilizes multiple mechanisms to counteract downstream inflammatory responses.

Importantly, our data demonstrated that CNTFRa is not only spared from protein-level downregulation but is also transcriptionally upregulated. CNTF signaling through CNTFRa–LIFR–gp130 is strongly neurotrophic and supports neuronal survival, an essential requirement for HSV-1 latency (91). This observation raised the possibility that the virus may benefit from this neuroprotective signaling. Indeed, we found that treatment of latently infected neuronal cells with recombinant CNTF induced viral reactivation from latency, identifying CNTF signaling as a previously underrecognized component of HSV-1 latency and reactivation. Supporting this hypothesis, clinical trials administering recombinant CNTF to amyotrophic lateral sclerosis patients (ALS) patients resulted in dose-dependent HSV-1 reactivation, suggesting CNTF signaling can modulate the balance between latency–reactivation (92). While ALS itself does not trigger HSV-1 reactivation (93), these clinical trial findings highlighted a potential mechanism by which CNTF signaling influences HSV-1 latency, likely by mimicking stress-related cytokine signaling (69). Previous studies identified IL-6 response elements within the HSV-1 genome, specifically concentrated in the promoter regions of both the latency-associated transcript (LAT) and immediate-early (IE) genes (94-97), indicating that IL-6 cytokines are a signal for virus reactivation for latency. Conversely, STAT3 has been shown to be required for the maintenance of the latent state of the virus (98).

In summary, our findings highlight the important role of gp130 receptor family signaling during both the lytic and latent stages of HSV-1 pathogenesis. To enable productive infection, the virus has evolved elegant strategies to downmodulate these receptors. Furthermore, while signaling molecules from these pathways are essential for maintaining latency, the activation of inflammatory signaling triggers viral reactivation.

## Materials and Methods

### Cell lines and viruses

The hTERT-HEL cells (immortalized human embryonic lung fibroblasts transformed with human telomerase reverse transcriptase [hTERT]) were cultured in Dulbecco’s modified Eagle’s medium (DMEM) supplemented with 10% fetal bovine serum (FBS). Vero, HEp-2, and LUHMES (Lund human mesencephalic) cells were obtained from the American Type Culture Collection (ATCC) and were cultured according to the manufacturer’s instructions. HSV-1(F) is a limited-passage isolate described before (99). The properties of the ΔICP0, ICP0 Δ244-277, and R2621 (ΔVHS) mutant viruses were described elsewhere (31, 100, 101).

### Western blot analysis

Cells were solubilized in triple detergent buffer (50 mM Tris-HCl [pH 8], 150 mM NaCl, 0.1% sodium dodecyl sulfate, 1% Nonidet P-40, 0.5% sodium deoxycholate, 100 mg/ml of phenylmethylsulfonyl fluoride) supplemented with phosphatase inhibitors (10 mM NaF, 10 mM β-glycerophosphate, 0.1 mM sodium vanadate) and protease inhibitor cocktail (Sigma), then briefly sonicated. Protein concentration was determined with Bradford (Bio-Rad Laboratories). The antibodies used and their dilutions are listed in the key resources table. Proteins were visualized with SuperSignal West Pico PLUS Chemiluminescent Substrate (Thermo Fisher Scientific) using the ImageQuant 800 western blot imaging system (Amersham).

### Real-time PCR analysis

Procedures for mRNA isolation, quantification, and real-time PCR were as described before (39, 102-104). Briefly, cells were lysed in 1 ml of TRIzol (Life Technologies), total RNA was extracted by phenol-chloroform, and followed by DNase treatment (Turbo DNase, Ambion). cDNA was generated from 1 ug of total RNA per sample using LunaScript® RT Master Mix Kit (Primer-free) (New England Biolabs E3025L). Real-time PCR analyses were performed using SYBR Green reagent (Invitrogen) or TaqMan (Applied Biosystems) according to the manufacturer’s recommendations in a QuantStudio 5 real-time PCR system (Applied Biosystems). Predesigned probes (6-carboxyfluorescein [FAM]/MGB) for the human *IL6ST, OSMRβ, LIFR, TNFR1, TNFR2, CNTFR, IL-31R, IL-6,* and *SOCS3* transcripts were obtained through Thermo Fisher Scientific and are listed in the Table 1. The 18S rRNA primers (universal primers; Ambion) were used for normalization. Primer pairs for the viral ICP27 and VP16 are listed in Table 1. Taqman probe for LAT is listed in Table 1.

**Table 1:**
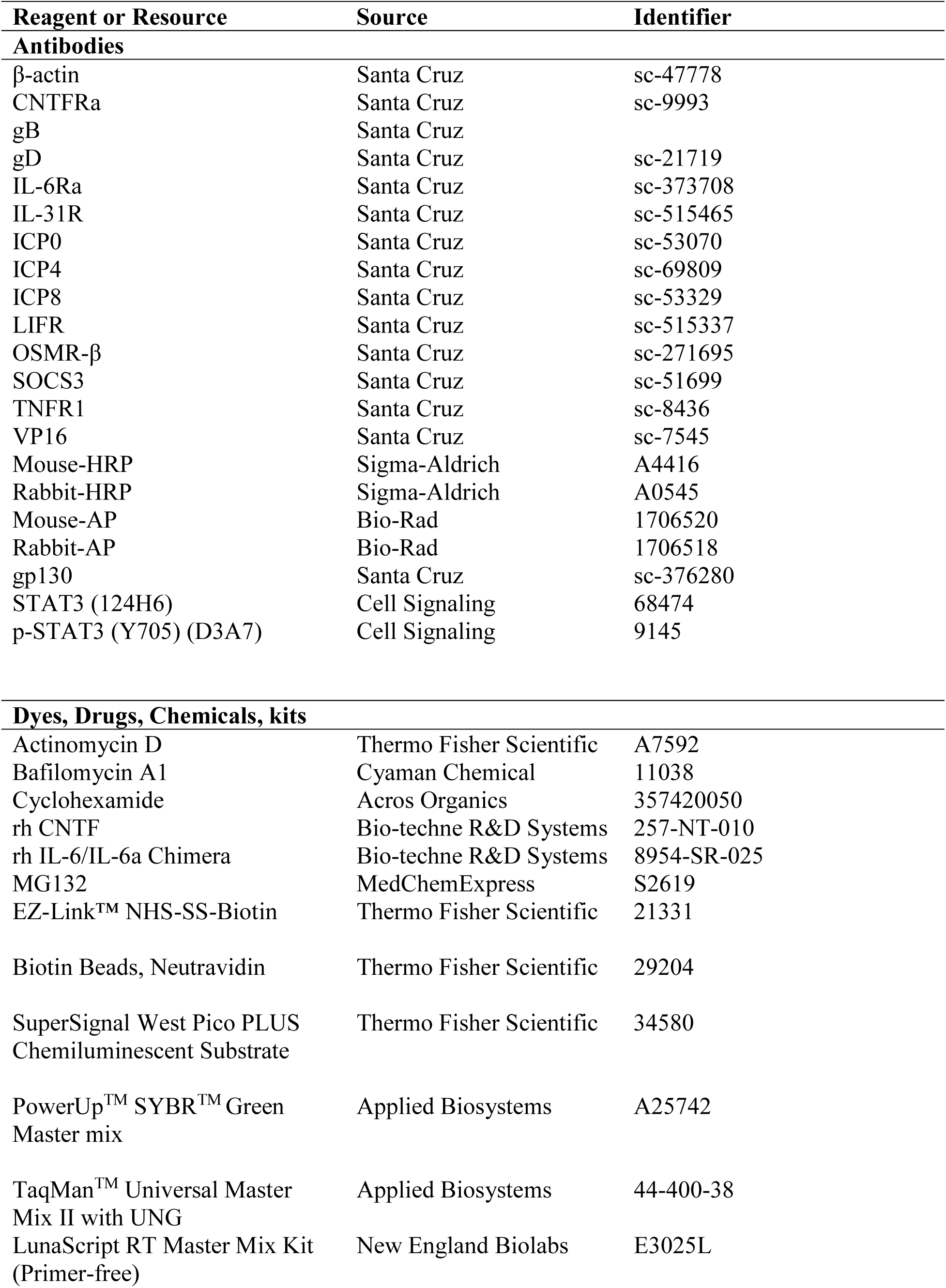
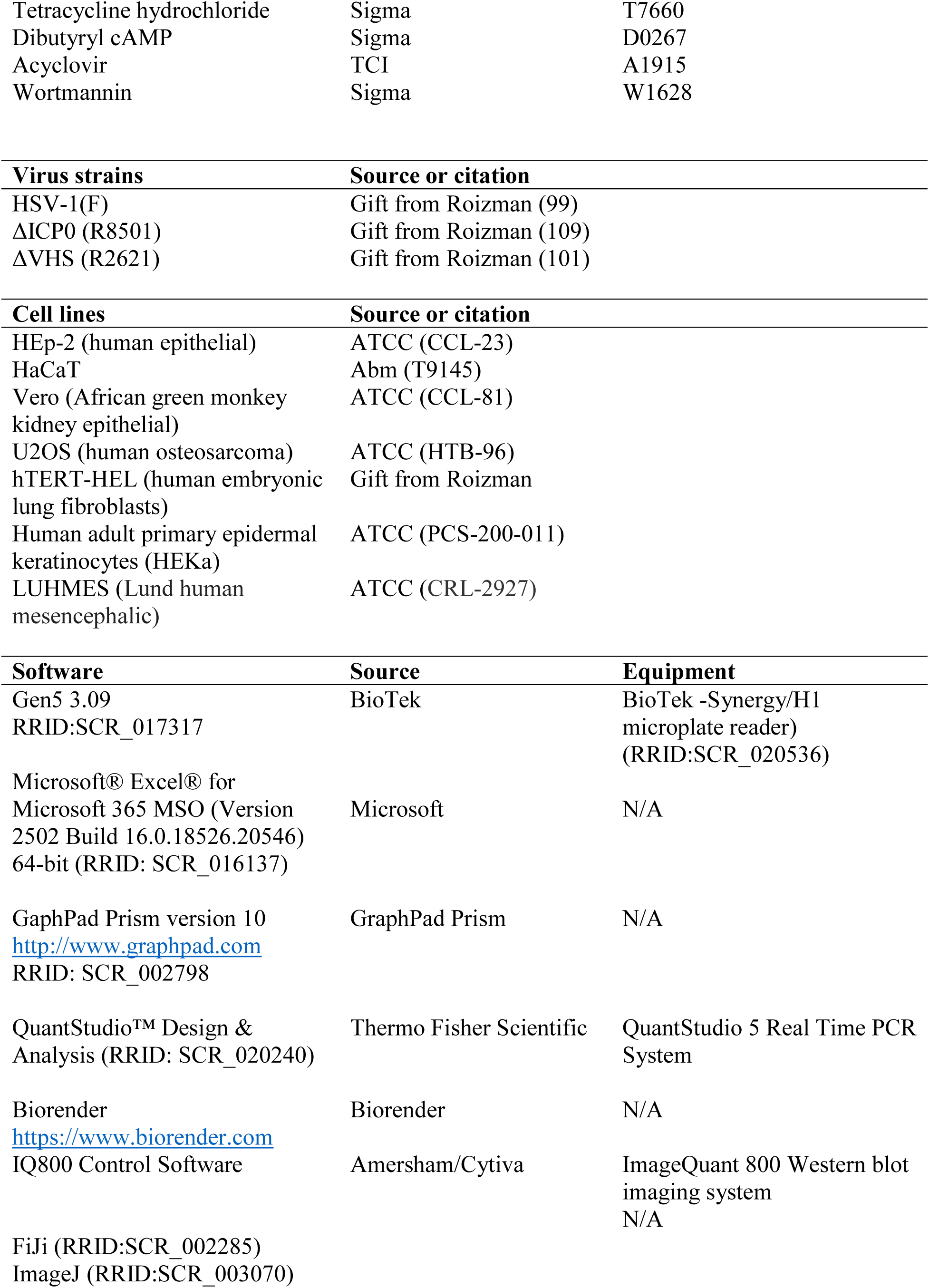
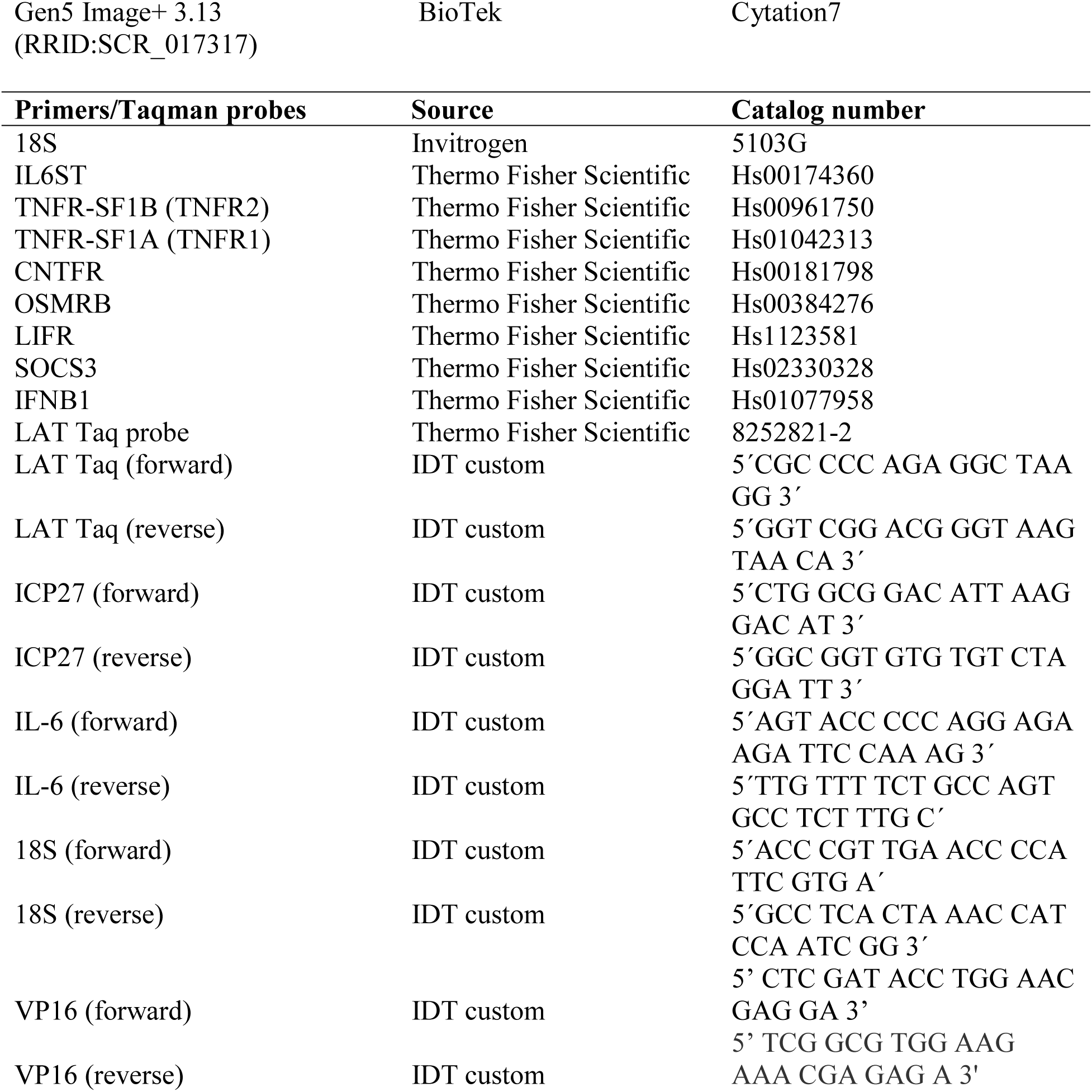
Key resources used in this study.

### Extracellular vesicle purification

Isolation of EVs from HSV-1(F)-infected or uninfected cells was done as previously described with minor modifications (39, 105, 106). Briefly, hTERT-HEL cells were infected with HSV-1(F) (0.5 PFU/cell). At 16 h postinfection, the supernatant was collected, centrifuged at 1,000x *g* for 10 min and at 2,000x *g* for 20 min to remove floating cells and larger cell debris, and filtered through a 0.45-μm-pore-size filter. The supernatant was concentrated (molecular mass cutoff, 30 kDa; Centricon Plus 70) according to the manufacturer’s instructions (Millipore). Samples were centrifuged in an SW41Ti rotor for 1.5 h at 100,000× *g* and 4°C in a Beckman Coulter Optima XPN-80 ultracentrifuge and pellets were collected for further analysis.

### HSV-1 latency establishment and reactivation in LUHMES

LUHMES cells were obtained from the ATCC and cultured in DMEM/F12 supplemented with N-2, as recommended by the supplier (107). Cells were grown in in matrigel-coated plates and allowed to proliferate up to 60-70% confluency, at which time the culture medium was changed to DMEM/F12 medium supplemented with differentiation factors, including: GDNF (recombinant human Glial cell Derived Neurotrophic Factor) (2 ng/ml), dibutyryl cAMP (N^6^,2′-O-Dibutyryladenosine 3′,5′-cyclic monophosphate sodium salt) (1 mM), and tetracycline (1.5ug/ml). Media was changed daily, and following 10 d of differentiation, postmitotic neurons were pretreated with 100 μM acyclovir (ACV) for 3 h and then infected with HSV-1 (F) (3 PFU/cell). ACV was removed at 48 hpi. The neurons were cultured to 21 dpi and replicates cultures (n=3) were harvested at 1, 3, 5, 9, 12, & 15 dpi to quantify LAT and ICP27 transcripts. At 19 dpi, rhCNTF (200 ng/ml), rhIL-6/Il-6Ra (200 ng/ml), or wortmannin (2 μM) were added to probe for reactivation. 48 h later the samples (n=3) were collected and analyzed for viral transcripts by RT-qPCR.

### Statistical analysis

The Prism 7 software (GraphPad) was used for statistical analysis of quantitative PCR (qPCR) data. *P* values were calculated using two-tailed Student’s *t* test. *P* values of < 0.05 were considered significant. Asterisks are defined as follows: P <0.05*; P < 0.01**; P < 0.001***; and P < 0.0001****. All statistical analyses were performed using at least three biological replicates.

### Cell surface biotinylation

hTERT-HEL were infected (10 PFU/cell) or left uninfected and surface labeling occurred at 12 hpi, as described before with minor modification (108). Briefly, cells (3 x 10^7^) were washed with ice-cold PBS, followed by incubation at 4^◦^C for 20 min to stop endocytosis. EZ-Link™ NHS-SS-Biotin (3mg total, 0.1 mg/ml) was added to the cultures, followed by additional incubation at 4^◦^C for 30 min. Cells were washed at 4^◦^C (3x) with 100 mM glycine in 10 min intervals on a shaker to quench the reactions and remove excess of biotin. Cells were detached by scraping and pelleted at 500x *g* for 10 min (4^◦^C). Cells were then lysed in 25 mM Tris-HCl pH 7.4, 150 mM NaCl, 1 mM EDTA, 1% NP-40, 5% glycerol, briefly sonicated, and centrifuged at 10,000x *g* for 1 min to remove larger debris. Neutravidin beads were added to the supernatant and incubated overnight on a rotator at 4^◦^C. Beads were then washed in the lysis buffer and bead-bound proteins were analyzed either by WB or LC-MS/MS.

### Mass Spectrometry Analysis (LC-MS/MS)

Proteins were eluted off agarose beads by washing 2x with 0.1 M glycine, pH 2, and samples were neutralized by adding 10 µl increments of 1 M TEAB (triethylammonium bicarbonate buffer pH 8.5) until pH was ∼7. Samples were dried in a speedvac and resuspended in 100 µl of 50 mM TEAB. Disulfides were reduced by adding 10 µl of 50 mM TCEP (5 mM final) and incubating at 55 ^0^C for 30 min. Cysteines were alkylated by adding 3µl of 375 mM IAA (iodoacetamide) (10 mM final) and incubating at RT in the dark for 30 min. Proteins were precipitated by adding 450 µl of acetone (1:5 dilution) and incubating at -20 ^0^C overnight. Samples were centrifuged at 14,000x *g* for 10 min at 4C. Supernatant was discarded, proteins were air dried, resuspended in 100 µl of 50 mM TEAB, 2 mM CaCl_2_, and digested by adding 500 ng of trypsin (0.1 mg/mL) overnight at 37 ^0^C. Reactions were quenched by the addition of 10% formic acid to 1%. Peptide concentrations were determined using the nanodrop UV system and based on the concentrations, 20 µl of each sample was loaded onto the C18 RP column and injected into the Orbitrap Ascend mass spectrometer with FAIMS for LC-MS/MS analysis. Data was searched using the Proteome Discoverer SeQuest software against the human database downloaded from UniProt on 05-05-2023 and HSV-1 database downloaded from UniProt on 05-15-2023.

### Analysis of proteomics data and heatmap construction

Normalized protein abundances from the surface proteomics experiment were used, including three HSV-1-infected replicates and three replicates from MOCK (uninfected) cells. Entries were restricted to Homo sapiens (OX=9606). Proteins lacking a gene symbol or abundance values in all six samples were removed.

For pathway analysis, proteins significantly downregulated or upregulated in HSV-1-infected cells relative to MOCK cells were identified. These proteins were submitted to the PANTHER classification system for Gene Ontology (GO) overrepresentation analysis using a reference list consisting of all proteins detected in the proteomics experiment. The enriched terms examined were membrane for downregulated proteins and cell surface for upregulated proteins. For each GO term, the PANTHER-derived list of UniProt accessions was cleaned and matched to the proteomic dataset by accession. Accessions not found in the quantified human protein dataset were excluded from visualization.

#### Replicate-based filtering and imputation of missing values

Abundances were log2-transformed because the surface proteomics data contained many missing values, which may reflect protein abundances below the limit of detection (LOD), such as after their removal from the cell surface. Proteins were first filtered based on the number of quantified replicates in each condition. Proteins detected in only one replicate in one condition and in no more than one replicate in the other condition (MOCK/HSV combinations of 1/1, 1/0, or 0/1) were removed because there was insufficient evidence for a reliable comparison.

A global LOD was defined as the 5^th^ percentile of all valid (non-missing) raw abundance values across the six samples and was then log2-transformed. Missing-value imputation was performed separately for each protein and within each condition (HSV-1 and MOCK), according to the number of replicates with measured values:

1. Three of three replicates quantified: No imputation was performed.
2. Two of three replicates quantified: The missing value was assumed to be missing at random and replaced with the mean of the two observed values for that condition.
3. One or zero of the three replicates quantified: The protein was considered absent or below the detection limit in that condition, and all three replicate values were set to a protein-specific floor. The floor was defined as the lower of (i) the global LOD and (ii) 0.1 log2 units below the lowest observed value for that protein across both conditions. This ensured that imputed values remained below all measured values for that protein.

#### Heatmap construction and hierarchical clustering

Imputed log2 abundances were standardized for each protein using a row-wise Z-score, with a mean of 0 and a standard deviation (SD) of 1 across the six samples. Thus, the heatmap represents relative abundance changes between mock and HSV-1 samples rather than absolute protein abundance. Proteins were hierarchically clustered using Euclidean distance with complete linkage. Columns were not clustered to preserve the experimental grouping. Heatmaps were generated in Python using seaborn.clustermap with a diverging color palette centered at zero. The color range was clipped to Z=-1.5 to +1.5 to maximize visual contrast. Every gene symbol was labeled according to the resulting clustered order, and proteins of biological interest were shown in bold.

## Acknowledgements

This study was supported by the NIAID R01AI162784 grant.

## REFERENCES

1. Knipe DM, Howley PM. 2007. Fields Virology Vol. 1 and 2.

2. Navarro AJ, Pasten IA, López-Hernández D, Bueno SM, Kalergis AM, González PA. 2026. The Expanding Organ Tropism of Herpes Simplex Virus Type 1. Rev Med Virol 36:e70155.

3. Saud R, Foster-Lemieur K, Duguay B, Swerdlow R, Kalamvoki M. 2026. Herpes simplex virus 1 subverts the mitochondrial network to support the infection: A lesson on mitochondrial versatility. bioRxiv doi:10.64898/2026.09.02.748825:2026.09.02.748825.

4. Nicoll MP, Proença JT, Efstathiou S. 2012. The molecular basis of herpes simplex virus latency. FEMS Microbiol Rev 36:684–705.

5. Dogrammatzis C, Waisner H, Kalamvoki M. 2020. “Non-Essential” Proteins of HSV-1 with Essential Roles In Vivo: A Comprehensive Review. Viruses 13.

6. Paludan SR. 2001. Requirements for the induction of interleukin-6 by herpes simplex virus-infected leukocytes. J Virol 75:8008–15.

7. Roca Suarez AA, Van Renne N, Baumert TF, Lupberger J. 2018. Viral manipulation of STAT3: Evade, exploit, and injure. PLoS Pathog 14:e1006839.

8. Li X, Huang C, Rai KR, Xu Q. 2025. Innate immune role of IL-6 in influenza a virus pathogenesis. Front Cell Infect Microbiol 15:1605446.

9. Osman EEA, Neamati N. 2024. Ironing Out the Mechanism of gp130 Signaling. Pharmacol Rev 76:1399–1443.

10. Kishimoto T. 1994. Signal transduction through homo- or heterodimers of gp130. Stem Cells 12 Suppl 1:37–44; discussion 44–5.

11. Silver JS, Hunter CA. 2010. gp130 at the nexus of inflammation, autoimmunity, and cancer. J Leukoc Biol 88:1145–56.

12. Rose-John S. 2012. IL-6 trans-signaling via the soluble IL-6 receptor: importance for the pro-inflammatory activities of IL-6. Int J Biol Sci 8:1237–47.

13. Shkhyan R, Flynn C, Lamoure E, Sarkar A, Van Handel B, Li J, York J, Banks N, Van der Horst R, Liu NQ, Lee S, Bajaj P, Vadivel K, Harn HI, Tassey J, Lozito T, Lieberman JR, Chuong CM, Hurtig MS, Evseenko D. 2023. Inhibition of a signaling modality within the gp130 receptor enhances tissue regeneration and mitigates osteoarthritis. Sci Transl Med 15:eabq2395.

14. Harker JA, Wong KA, Dolgoter A, Zuniga EI. 2015. Cell-Intrinsic gp130 Signaling on CD4+ T Cells Shapes Long-Lasting Antiviral Immunity. J Immunol 195:1071–81.

15. Gu R, Zhang H, Xu E, Liu S. 2025. The regulatory roles of STAT3 protein in the pathogenesis of viral infections. Front Cell Infect Microbiol 15:1706270.

16. Ho HH, Ivashkiv LB. 2006. Role of STAT3 in type I interferon responses. Negative regulation of STAT1-dependent inflammatory gene activation. J Biol Chem 281:14111–8.

17. Suthaus J, Adam N, Grötzinger J, Scheller J, Rose-John S. 2011. Viral Interleukin-6: Structure, pathophysiology and strategies of neutralization. Eur J Cell Biol 90:495–504.

18. Komaki S, Inagaki T, Kumar A, Izumiya Y. 2024. The Role of vIL-6 in KSHV-Mediated Immune Evasion and Tumorigenesis. Viruses 16.

19. Qiu Q, Yang X, Li X, Ren Y, Huang N. 2025. Viral IL-10 can promote the proliferation, migration and invasion of nasopharyngeal carcinoma cells and inhibit their apoptosis. Discov Oncol 16:199.

20. Ren Y, Yang J, Li M, Huang N, Chen Y, Wu X, Wang X, Qiu S, Wang H, Li X. 2020. Viral IL-10 promotes cell proliferation and cell cycle progression via JAK2/STAT3 signaling pathway in nasopharyngeal carcinoma cells. Biotechnol Appl Biochem 67:929– 938.

21. Slinger E, Maussang D, Schreiber A, Siderius M, Rahbar A, Fraile-Ramos A, Lira SA, Söderberg-Nauclér C, Smit MJ. 2010. HCMV-encoded chemokine receptor US28 mediates proliferative signaling through the IL-6-STAT3 axis. Sci Signal 3:ra58.

22. Schuster B, Kovaleva M, Sun Y, Regenhard P, Matthews V, Grötzinger J, Rose-John S, Kallen K-J. 2003. Signaling of Human Ciliary Neurotrophic Factor (CNTF) Revisited: THE INTERLEUKIN-6 RECEPTOR CAN SERVE AS AN α-RECEPTOR FOR CNTF *. Journal of Biological Chemistry 278:9528–9535.

23. Ramón Roth I, Führing JI, Garbers C. 2026. Understanding complex formation of gp130 cytokines for the design of selective therapeutics. Biochim Biophys Acta Mol Cell Res 1873:120105.

24. Bravo J, Heath JK. 2000. Receptor recognition by gp130 cytokines. Embo j 19:2399–411.

25. Zhu H, Zheng C. 2020. The Race between Host Antiviral Innate Immunity and the Immune Evasion Strategies of Herpes Simplex Virus 1. Microbiol Mol Biol Rev 84.

26. Su D, Han L, Shi C, Li Y, Qian S, Feng Z, Yu L. 2024. An updated review of HSV-1 infection-associated diseases and treatment, vaccine development, and vector therapy application. Virulence 15:2425744.

27. Smith JB, Herbert JJ, Truong NR, Cunningham AL. 2022. Cytokines and chemokines: The vital role they play in herpes simplex virus mucosal immunology. Frontiers in Immunology Volume 13–2022.

28. Birzer A, Krawczyk A, Draßner C, Kuhnt C, Mühl-Zürbes P, Heilingloh CS, Steinkasserer A, Popella L. 2020. HSV-1 Modulates IL-6 Receptor Expression on Human Dendritic Cells. Front Immunol 11:1970.

29. Liang L, Roizman B. 2006. Herpes simplex virus 1 precludes replenishment of the short-lived receptor of tumor necrosis factor alpha by virion host shutoff-dependent degradation of its mRNA. J Virol 80:7756–9.

30. Waisner H, Kalamvoki M. 2019. The ICP0 Protein of Herpes Simplex Virus 1 (HSV-1) Downregulates Major Autophagy Adaptor Proteins Sequestosome 1 and Optineurin during the Early Stages of HSV-1 Infection. J Virol 93.

31. Waisner H, Saud R, Lasnier S, Suma SM, Foster-Lemieur K, Kalamvoki M. 2025. Herpes simplex virus diverts CIN85 endosomal cargo for exocytosis to evade antiviral responses: a novel role for the viral immediate-early protein ICP0. mBio 16:e0214325.

32. Deschamps T, Dogrammatzis C, Mullick R, Kalamvoki M. 2017. Cbl E3 Ligase Mediates the Removal of Nectin-1 from the Surface of Herpes Simplex Virus 1-Infected Cells. Journal of Virology 91:10.1128/jvi.00393–17.

33. Liang Y, Kurakin A, Roizman B. 2005. Herpes simplex virus 1 infected cell protein 0 forms a complex with CIN85 and Cbl and mediates the degradation of EGF receptor from cell surfaces. Proc Natl Acad Sci U S A 102:5838–43.

34. Lasnier S, Kalamvoki M. 2026. Effect of HSV-1 ICP0 E3 ubiquitin ligase on CIN85 endosomes. bioRxiv doi:10.64898/2026.09.21.753399:2026.09.21.753399.

35. Kwong AD, Frenkel N. 1987. Herpes simplex virus-infected cells contain a function(s) that destabilizes both host and viral mRNAs. Proceedings of the National Academy of Sciences 84:1926–1930.

36. Smiley JR. 2004. Herpes simplex virus virion host shutoff protein: immune evasion mediated by a viral RNase? J Virol 78:1063–8.

37. Dogrammatzis C, Saud R, Waisner H, Lasnier S, Suma SM, Grieshaber B, Kalamvoki M. 2024. Tracing the STING exocytosis pathway during herpes viruses infection. mBio 15:e0037324.

38. Waisner H, Lasnier S, Suma SM, Kalamvoki M. 2023. Effects on exocytosis by two HSV-1 mutants unable to block autophagy. J Virol 97:e0075723.

39. Dogrammatzis C, Saleh S, Deighan C, Kalamvoki M. 2021. Diverse Populations of Extracellular Vesicles with Opposite Functions during Herpes Simplex Virus 1 Infection. J Virol 95.

40. Wang WB, Levy DE, Lee CK. 2011. STAT3 negatively regulates type I IFN-mediated antiviral response. J Immunol 187:2578–85.

41. Chang Z, Wang Y, Zhou X, Long JE. 2018. STAT3 roles in viral infection: antiviral or proviral? Future Virol 13:557–574.

42. Crow MS, Lum KK, Sheng X, Song B, Cristea IM. 2016. Diverse mechanisms evolved by DNA viruses to inhibit early host defenses. Crit Rev Biochem Mol Biol 51:452–481.

43. Croker BA, Krebs DL, Zhang JG, Wormald S, Willson TA, Stanley EG, Robb L, Greenhalgh CJ, Förster I, Clausen BE, Nicola NA, Metcalf D, Hilton DJ, Roberts AW, Alexander WS. 2003. SOCS3 negatively regulates IL-6 signaling in vivo. Nat Immunol 4:540–5.

44. Baker BJ, Akhtar LN, Benveniste EN. 2009. SOCS1 and SOCS3 in the control of CNS immunity. Trends Immunol 30:392–400.

45. Alston CI, Dix RD. 2019. SOCS and Herpesviruses, With Emphasis on Cytomegalovirus Retinitis. Front Immunol 10:732.

46. Rock DL, Fraser NW. 1983. Detection of HSV-1 genome in central nervous system of latently infected mice. Nature 302:523–5.

47. Roizman B, Whitley RJ. 2013. An inquiry into the molecular basis of HSV latency and reactivation. Annu Rev Microbiol 67:355–74.

48. Stevens JG, Wagner EK, Devi-Rao GB, Cook ML, Feldman LT. 1987. RNA complementary to a herpesvirus alpha gene mRNA is prominent in latently infected neurons. Science 235:1056–9.

49. Camarena V, Kobayashi M, Kim JY, Roehm P, Perez R, Gardner J, Wilson AC, Mohr I, Chao MV. 2010. Nature and duration of growth factor signaling through receptor tyrosine kinases regulates HSV-1 latency in neurons. Cell Host Microbe 8:320–30.

50. Kim JY, Mandarino A, Chao MV, Mohr I, Wilson AC. 2012. Transient reversal of episome silencing precedes VP16-dependent transcription during reactivation of latent HSV-1 in neurons. PLoS Pathog 8:e1002540.

51. Laycock KA, Lee SF, Brady RH, Pepose JS. 1991. Characterization of a murine model of recurrent herpes simplex viral keratitis induced by ultraviolet B radiation. Invest Ophthalmol Vis Sci 32:2741–6.

52. Suzich JB, Cliffe AR. 2018. Strength in diversity: Understanding the pathways to herpes simplex virus reactivation. Virology 522:81–91.

53. Edwards TG, Bloom DC. 2019. Lund Human Mesencephalic (LUHMES) Neuronal Cell Line Supports Herpes Simplex Virus 1 Latency In Vitro. J Virol 93.

54. Liu X, Cohen JI. 2015. The role of PI3K/Akt in human herpesvirus infection: From the bench to the bedside. Virology 479-480:568–77.

55. Nicola AV, Straus SE. 2004. Cellular and viral requirements for rapid endocytic entry of herpes simplex virus. J Virol 78:7508–17.

56. Huang GN. 2012. Biotinylation of Cell Surface Proteins. Bio Protoc 2.

57. Shi Y, Zhou W, Cheng L, Chen C, Huang Z, Fang X, Wu Q, He Z, Xu S, Lathia JD, Ping Y, Rich JN, Bian XW, Bao S. 2017. Tetraspanin CD9 stabilizes gp130 by preventing its ubiquitin-dependent lysosomal degradation to promote STAT3 activation in glioma stem cells. Cell Death Differ 24:167–180.

58. Schumacher N, Thomsen I, Brundert F, Hejret V, Düsterhöft S, Tichý B, Schmidt-Arras D, Voss M, Rose-John S. 2023. EGFR stimulation enables IL-6 trans-signalling via iRhom2-dependent ADAM17 activation in mammary epithelial cells. Biochim Biophys Acta Mol Cell Res 1870:119489.

59. Calligaris M, Yang CY, Bonelli S, Spanò DP, Müller SA, Lichtenthaler SF, Troeberg L, Scilabra SD. 2023. Identification of membrane proteins regulated by ADAM15 by SUSPECS proteomics. Front Mol Biosci 10:1162504.

60. Itagaki M, Kamei N. 2025. Clathrin functions in intracellular sorting during receptor-mediated endocytosis. FEBS Lett 599:1509–1517.

61. Collins MV, Lång HKM, Finkielstein CV, Ryhänen SJ, Ikonen E, Capelluto DGS. 2026. TOM1 family proteins: from cargo sorting to immune dysregulation and cancer. Cell Commun Signal 24.

62. Goldenring JR. 2015. Recycling endosomes. Curr Opin Cell Biol 35:117–22.

63. Puthenveedu MA, Lauffer B, Temkin P, Vistein R, Carlton P, Thorn K, Taunton J, Weiner OD, Parton RG, von Zastrow M. 2010. Sequence-dependent sorting of recycling proteins by actin-stabilized endosomal microdomains. Cell 143:761–73.

64. Bögershausen N, Cavdarli B, Nagai TH, Milev MP, Wolff A, Mehranfar M, Schmidt J, Choudhary D, Gutiérrez-Gutiérrez Ó, Cyganek L, Saint-Dic D, Zibat A, Köhrer K, Wollenweber TE, Wieczorek D, Altmüller J, Borodina T, Kaçar D, Haliloğlu G, Li Y, Thiel C, Sacher M, Knapik EW, Yigit G, Wollnik B. 2025. SEC24C deficiency causes trafficking and glycosylation abnormalities in an epileptic encephalopathy with cataracts and dyserythropoeisis. JCI Insight 10.

65. Asai M, Hattori C, Szabó B, Sasagawa N, Maruyama K, Tanuma S, Ishiura S. 2003. Putative function of ADAM9, ADAM10, and ADAM17 as APP alpha-secretase. Biochem Biophys Res Commun 301:231–5.

66. Zobiack N, Rescher U, Ludwig C, Zeuschner D, Gerke V. 2003. The annexin 2/S100A10 complex controls the distribution of transferrin receptor-containing recycling endosomes. Mol Biol Cell 14:4896–908.

67. Tanaka T, Narazaki M, Kishimoto T. 2014. IL-6 in inflammation, immunity, and disease. Cold Spring Harb Perspect Biol 6:a016295.

68. Heinrich PC, Behrmann I, Haan S, Hermanns HM, Müller-Newen G, Schaper F. 2003. Principles of interleukin (IL)-6-type cytokine signalling and its regulation. Biochem J 374:1–20.

69. Riethmueller S, Ehlers JC, Lokau J, Düsterhöft S, Knittler K, Dombrowsky G, Grötzinger J, Rabe B, Rose-John S, Garbers C. 2016. Cleavage Site Localization Differentially Controls Interleukin-6 Receptor Proteolysis by ADAM10 and ADAM17. Sci Rep 6:25550.

70. Schumacher N, Rose-John S. 2019. ADAM17 Activity and IL-6 Trans-Signaling in Inflammation and Cancer. Cancers (Basel) 11.

71. Rose-John S, Jenkins BJ, Garbers C, Moll JM, Scheller J. 2023. Targeting IL-6 trans-signalling: past, present and future prospects. Nat Rev Immunol 23:666–681.

72. Taga T, Hibi M, Hirata Y, Yamasaki K, Yasukawa K, Matsuda T, Hirano T, Kishimoto T. 1989. Interleukin-6 triggers the association of its receptor with a possible signal transducer, gp130. Cell 58:573–81.

73. Taga T, Kishimoto T. 1997. Gp130 and the interleukin-6 family of cytokines. Annu Rev Immunol 15:797–819.

74. Kishimoto T, Akira S, Narazaki M, Taga T. 1995. Interleukin-6 family of cytokines and gp130. Blood 86:1243–54.

75. Jostock T, Müllberg J, Ozbek S, Atreya R, Blinn G, Voltz N, Fischer M, Neurath MF, Rose-John S. 2001. Soluble gp130 is the natural inhibitor of soluble interleukin-6 receptor transsignaling responses. Eur J Biochem 268:160–7.

76. Baran P, Hansen S, Waetzig GH, Akbarzadeh M, Lamertz L, Huber HJ, Ahmadian MR, Moll JM, Scheller J. 2018. The balance of interleukin (IL)-6, IL-6·soluble IL-6 receptor (sIL-6R), and IL-6·sIL-6R·sgp130 complexes allows simultaneous classic and trans-signaling. J Biol Chem 293:6762–6775.

77. Gaucherand L, Gaglia MM. 2022. The Role of Viral RNA Degrading Factors in Shutoff of Host Gene Expression. Annu Rev Virol 9:213–238.

78. García-Sastre A. 2017. Ten Strategies of Interferon Evasion by Viruses. Cell Host Microbe 22:176–184.

79. Hsia HC, Stopford CM, Zhang Z, Damania B, Baldwin AS. 2017. Signal transducer and activator of transcription 3 (Stat3) regulates host defense and protects mice against herpes simplex virus-1 (HSV-1) infection. J Leukoc Biol 101:1053–1064.

80. Harrison Angela R, Moseley Gregory W. 2020. The Dynamic Interface of Viruses with STATs. Journal of Virology 94:10.1128/jvi.00856–20.

81. Molden J, Chang Y, You Y, Moore PS, Goldsmith MA. 1997. A Kaposi’s sarcoma-associated herpesvirus-encoded cytokine homolog (vIL-6) activates signaling through the shared gp130 receptor subunit. J Biol Chem 272:19625–31.

82. Sakakibara S, Tosato G. 2011. Viral interleukin-6: role in Kaposi’s sarcoma-associated herpesvirus: associated malignancies. J Interferon Cytokine Res 31:791–801.

83. Bermudez Y, Schultz S, Miles J, Dwyer W, Muller M. 2025. Virally encoded interleukin-6 (vIL-6) coordinates with human IL-6 to modulate cytokine expression during KSHV infection. bioRxiv doi:10.1101/2025.10.02.679950.

84. Adrain C, Zettl M, Christova Y, Taylor N, Freeman M. 2012. Tumor necrosis factor signaling requires iRhom2 to promote trafficking and activation of TACE. Science 335:225–8.

85. Al-Salihi MA, Lang PA. 2020. iRhom2: An Emerging Adaptor Regulating Immunity and Disease. Int J Mol Sci 21.

86. Wolf J, Waetzig GH, Chalaris A, Reinheimer TM, Wege H, Rose-John S, Garbers C. 2016. Different Soluble Forms of the Interleukin-6 Family Signal Transducer gp130 Fine-tune the Blockade of Interleukin-6 Trans-signaling. J Biol Chem 291:16186–96.

87. Schumertl T, Lokau J, Rose-John S, Garbers C. 2022. Function and proteolytic generation of the soluble interleukin-6 receptor in health and disease. Biochim Biophys Acta Mol Cell Res 1869:119143.

88. Noda-Nicolau NM, Polettini J, da Silva MG, Peltier MR, Menon R. 2018. Polybacterial stimulation suggests discrete IL-6/IL-6R signaling in human fetal membranes: Potential implications on IL-6 bioactivity. J Reprod Immunol 126:60–68.

89. Lemmers A, Gustot T, Durnez A, Evrard S, Moreno C, Quertinmont E, Vercruysse V, Demetter P, Franchimont D, Le Moine O, Geerts A, Devière J. 2009. An inhibitor of interleukin-6 trans-signalling, sgp130, contributes to impaired acute phase response in human chronic liver disease. Clin Exp Immunol 156:518–27.

90. Dominguez E, Mauborgne A, Mallet J, Desclaux M, Pohl M. 2010. SOCS3-mediated blockade of JAK/STAT3 signaling pathway reveals its major contribution to spinal cord neuroinflammation and mechanical allodynia after peripheral nerve injury. J Neurosci 30:5754–66.

91. Schuster B, Kovaleva M, Sun Y, Regenhard P, Matthews V, Grötzinger J, Rose-John S, Kallen KJ. 2003. Signaling of human ciliary neurotrophic factor (CNTF) revisited. The interleukin-6 receptor can serve as an alpha-receptor for CTNF. J Biol Chem 278:9528–35.

92. Miller RG, Bryan WW, Dietz MA, Munsat TL, Petajan JH, Smith SA, Goodpasture JC. 1996. Toxicity and tolerability of recombinant human ciliary neurotrophic factor in patients with amyotrophic lateral sclerosis. Neurology 47:1329–31.

93. Zheng Q, Wang D, Lin R, Chen Y, Huang H, Xu Z, Zheng C, Xu W. 2023. Mendelian randomization analysis suggests no associations of human herpes viruses with amyotrophic lateral sclerosis. Front Neurosci 17:1299122.

94. Kriesel JD, Ricigliano J, Spruance SL, Garza HH, Jr., Hill JM. 1997. Neuronal reactivation of herpes simplex virus may involve interleukin-6. J Neurovirol 3:441–8.

95. Al-Dujaili LJ, Clerkin PP, Clement C, McFerrin HE, Bhattacharjee PS, Varnell ED, Kaufman HE, Hill JM. 2011. Ocular herpes simplex virus: how are latency, reactivation, recurrent disease and therapy interrelated? Future Microbiol 6:877–907.

96. Duan F, Liao J, Huang Q, Nie Y, Wu K. 2012. HSV-1 miR-H6 inhibits HSV-1 replication and IL-6 expression in human corneal epithelial cells in vitro. Clin Dev Immunol 2012:192791.

97. Gessa-Sorroche M, Kanclerz P, Alio J. 2022. Evidence in the prevention of the recurrence of herpes simplex and herpes zoster keratitis after eye surgery. Arch Soc Esp Oftalmol (Engl Ed) 97:149–160.

98. Du T, Zhou G, Roizman B. 2013. Modulation of reactivation of latent herpes simplex virus 1 in ganglionic organ cultures by p300/CBP and STAT3. Proc Natl Acad Sci U S A 110:E2621–8.

99. Ejercito PM, Kieff ED, Roizman B. 1968. Characterization of herpes simplex virus strains differing in their effects on social behaviour of infected cells. J Gen Virol 2:357–64.

100. Horsburgh BC, Hubinette MM, Tufaro F. 1999. Genetic manipulation of herpes simplex virus using bacterial artificial chromosomes. Methods Enzymol 306:337–52.

101. Poon AP, Roizman B. 1997. Differentiation of the shutoff of protein synthesis by virion host shutoff and mutant gamma (1)34.5 genes of herpes simplex virus 1. Virology 229:98– 105.

102. Deschamps T, Kalamvoki M. 2017. Evasion of the STING DNA-Sensing Pathway by VP11/12 of Herpes Simplex Virus 1. J Virol 91.

103. Kalamvoki M, Roizman B. 2010. Interwoven Roles of Cyclin D3 and cdk4 Recruited by ICP0 and ICP4 in the Expression of Herpes Simplex Virus Genes. Journal of Virology 84:9709–9717.

104. Deschamps T, Kalamvoki M. 2017. Impaired STING Pathway in Human Osteosarcoma U2OS Cells Contributes to the Growth of ICP0-Null Mutant Herpes Simplex Virus. Journal of Virology 91:10.1128/jvi.00006–17.

105. Deschamps T, Kalamvoki M. 2018. Extracellular Vesicles Released by Herpes Simplex Virus 1-Infected Cells Block Virus Replication in Recipient Cells in a STING-Dependent Manner. J Virol 92.

106. Kalamvoki M. 2025. HSV-1 virions and related particles: biogenesis and implications in the infection. J Virol 99:e0107624.

107. Scholz D, Pöltl D, Genewsky A, Weng M, Waldmann T, Schildknecht S, Leist M. 2011. Rapid, complete and large-scale generation of post-mitotic neurons from the human LUHMES cell line. J Neurochem 119:957–71.

108. Deschamps T, Dogrammatzis C, Mullick R, Kalamvoki M. 2017. Cbl E3 Ligase Mediates the Removal of Nectin-1 from the Surface of Herpes Simplex Virus 1-Infected Cells. J Virol 91.

109. Gu H, Roizman B. 2007. Herpes simplex virus-infected cell protein 0 blocks the silencing of viral DNA by dissociating histone deacetylases from the CoREST-REST complex. Proc Natl Acad Sci U S A 104:17134–9.

